# Corpus Colossal: A Bibliometric Analysis of Neuroscience Abstracts and Impact Factor

**DOI:** 10.1101/505057

**Authors:** William M. Kenkel

**Affiliations:** Neuroscience Institute, Georgia State University

## Abstract

A field’s priorities are thought to be reflected by the contents of its high-impact journals. Researchers in turn may choose to pursue research objectives based on what is believed to be most highly valued by their peers. By compiling a corpus of abstracts from within the field neuroscience, I was able to analyze which terms had differential frequencies between 12 high-impact and 13 medium-impact journals. Approximately 50,000 neuroscience abstracts were analyzed over the years 2014-2018. Several broad trends emerged from the analysis of which terms were biased towards high-impact journals. Generally speaking, high-impact journals tended to feature: genetic or psychiatric studies, use of the latest and most sophisticated methods, examinations of the orbitofrontal cortex or amygdala, and/or use of human or non-mammalian subjects. Medium-impact journals tended to feature motor or cardiovascular studies, use of older methods, examinations of caudal brain regions, and/or rats as subjects.

## INTRODUCTION

A journal’s impact factor is determined by the number of citations received relative to the number of articles published. Within the culture of academic research, the contents of high-impact journals are therefore taken as a proxy for the interests and priorities of the field. Researchers use their sense of the field’s priorities to dictate their own research decisions, as well as in evaluating others’ work. These impressions are shaped by experience, training, and social conditioning, often without much systematic analysis.

Previous efforts have identified the 100 most-cited papers in neuroscience [1] or identified factors associated with high impact, such as age [2] or gender [3-5]. Among high-impact neuroscience and multidisciplinary journals, women authors are persistently underrepresented [3]. Scholars from younger generations receive less recognition despite publishing in better journals [2]. Across a researcher’s career, the number of citations peak in the early years then continually decline, while the impact factor of journals they publish in remains fairly stable [2]. These patterns are important to understand because there is generally a Matthew effect acting on research careers [2, 6, 7], such that early advantages accumulate and increase the likelihood of access to further advantage.

The most complete analysis of neuroscientific publication trends analyzed work from 2006-2015 [8]. This work, by Yeung et al., analyzed the patterns in citations among individual articles and observed a shift of focus from general brain imaging terms to cellular, molecular and genetic terms over the study period. The purpose of the present work was to take a bird’s-eye view of recent neuroscience publications and determine patterns in the terms that are differentially used in high-vs. medium-impact journals.

## METHODS

Abstracts were gathered from 12 high-impact and 13 medium-impact journals, creating two corpora spanning January 2014 to December 2018. Journals were selected according to the following criteria: 1) each journal must have a broad scope within the field of neuroscience, that is, they could not be limited to a single sub-discipline, method, or species; and 2) journals must follow conventional practices for abstracts and issue composition. Thus, in addition to many journals being excluded for being too narrowly focused, three high-impact journals in particular were left out: *Brain and Behavioral Sciences* was excluded for being too theoretical and dissimilar to other journals’ content; *Trends in Neuroscience* was excluded for having too brief abstracts; and *Progress in Brain Research* was excluded for having nonconventional issue composition. For three high-impact journals with especially broad, transdisciplinary scopes (*Science, Nature,* and *Nature Communications*) a further criterion was applied such that each abstract was scanned for occurrences of a neuroscience-related keyword and only included if at least one such keyword was present. The list of high- and medium-impact journals is shown in Table 1; the list of keywords indicated neuroscience relevance within the three broad scope high-impact journals is shown in Table 2. Truly low-impact journals were avoided as being overly niche. For medium-impact journals, impact factor ranged from 1.26 – 4.23, with a median of 2.83. For high-impact journals, impact factor ranged from 10.85 – 41.58, with a median of 14.5.

**Table 1.**
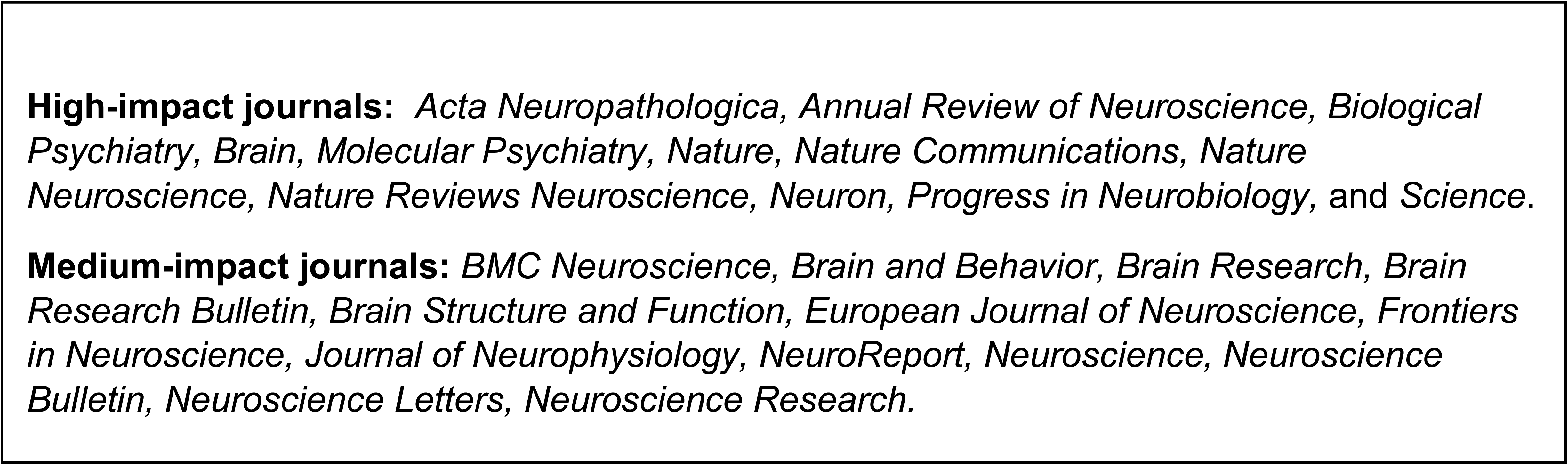

**Table 2.**
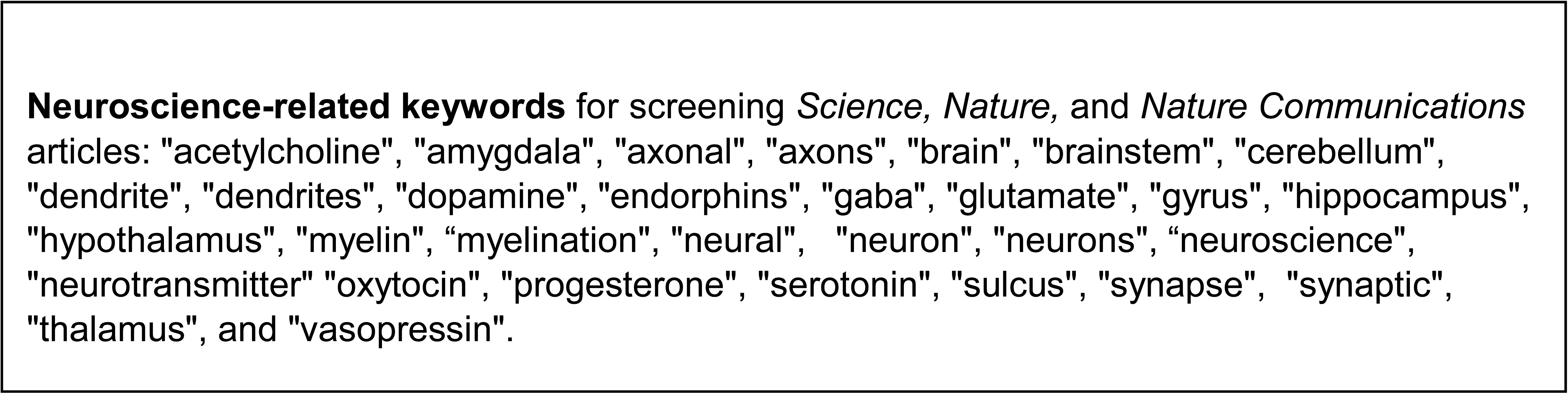

Abstracts were harvested from PubMed using the *PubMedWordCloud* package for R. All text was then converted to lower case, but for the sake of clarity will be shown in its most-common form in the following text and legends. After removing numbers, punctuation, and commonly used English stop words (e.g. ‘a’, ‘is’, ‘the’), each corpus was cleaned of generic research terms (e.g. ‘effect’, ‘group’, ‘increased’). A further set of terms was then removed for being either spurious or accidents of publication (e.g. ‘copyright’, ‘Ireland’, ‘university’). Various typographical errors were addressed on an as-needed basis, which consisted chiefly of incorrectly conjoined words (e.g. ‘patientderived’), reconciling certain plurals (e.g. combining ‘tumour’ and ‘tumours’), or resolving discrepant spellings between American and British English.

Each corpus was then analyzed using the *tidytext* and *tm* (text mining) packages for R. The R scripts used for analysis and raw data are attached as supplementary files. Incidences of each unique term were calculated and then compared between high-and medium-impact journals, controlling for each corpus’ overall word count. Each term was then assigned a log odds ratio derived from the ratio of occurrences in medium-vs. high-impact journals. Thus, a higher log odds ratio indicates a given term to be favored in high-impact journal abstracts. In order to be included in the final analyses, a term had to be used more than 10 times per year.

Secondary analyses were carried out comparing 1) study organisms, 2) brain regions, 3) neurotransmitters, 4) methodological approaches, and 5) broad themes within neuroscience, comparing in each case the log odds ratio of occurrence in medium-vs. high-impact journals. Each broad theme from within the larger field of neuroscience was assigned several keywords intended to be specific to that particular theme, as shown in Table 3. Each methodological approach was first evaluated for the specific term that was most frequently used (e.g. ‘optogenetics’ rather than ‘optogenetically’). The various conjugations of these methodological terms did not meaningfully differ in terms of odds ratio of occurrence.

**Table 3.**
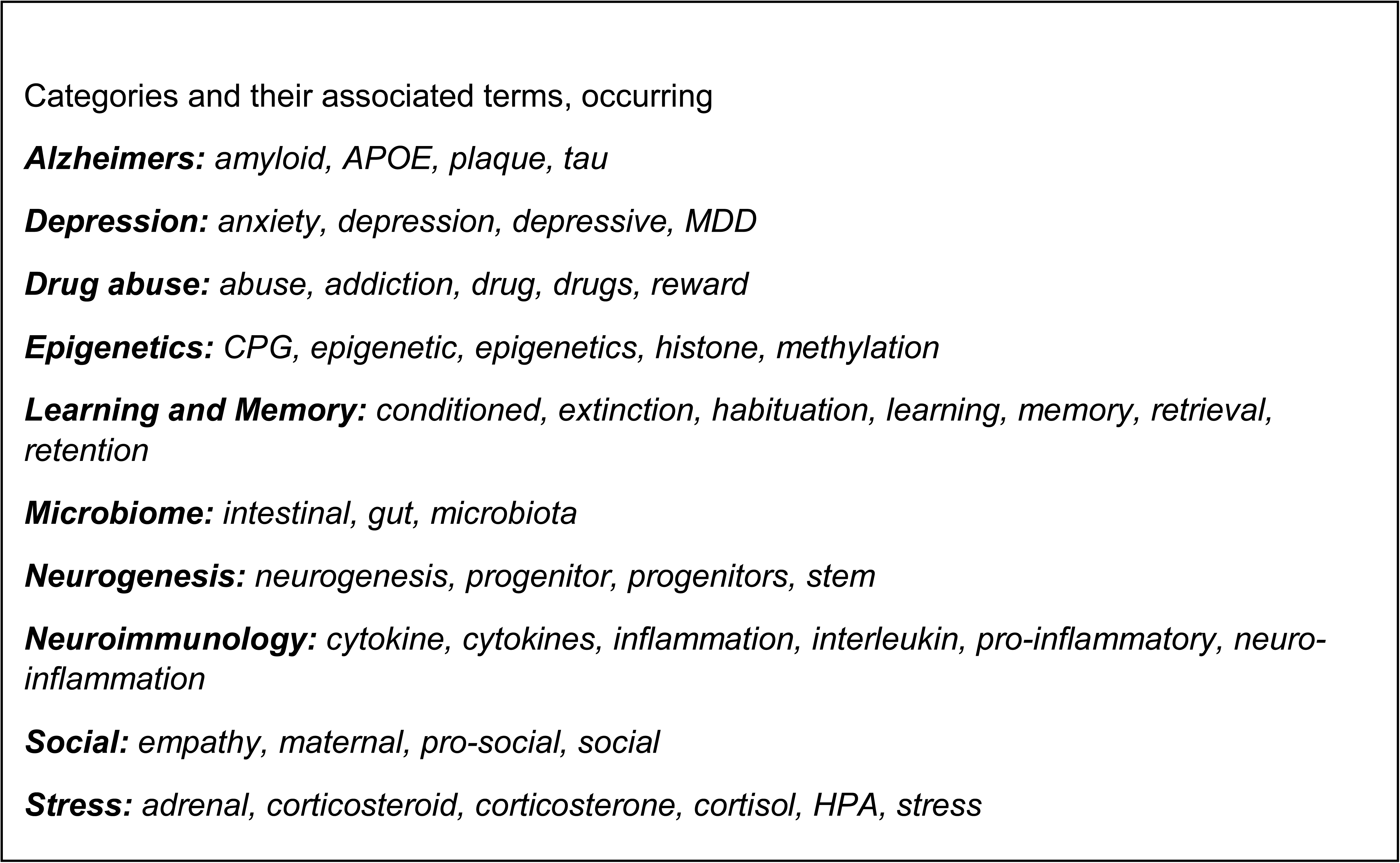

## RESULTS

A total of 13,059 abstracts were gathered from high-impact journals and 36,787 abstracts from medium-impact journals. The distribution of impact factor scores among the two categories of journals is shown in Figure 1A. The distribution of terms’ log odd ratios was roughly symmetrical, with slightly more terms differentially preferred by medium-impact journals (Figure 1B). On a year-by-year basis, terms scoring in the top 15 most differentiated (biased either towards high- or medium-impact journals) are shown in Figure 2. Over the entirety of the 5-year period, the terms scoring in the top 25 most differentiated were categorized into themes as shown in Figure 3. Secondary analyses were conducted across the 5-year study period, comparing study organisms (Figure 4), brain regions (Figure 5), neurotransmitters (Figure 6), methodological approaches (Figure 7), and broad themes within neuroscience (Figure 8). In the cases of study organism, neurotransmitter and approach, the size of each dot represents the total number of instances each term occurred. For instance, rats were found to be commonly used and associated with medium-impact journals, while *C. elegans* was found to be uncommonly used and associated with high-impact journals.

**Figure 1.**
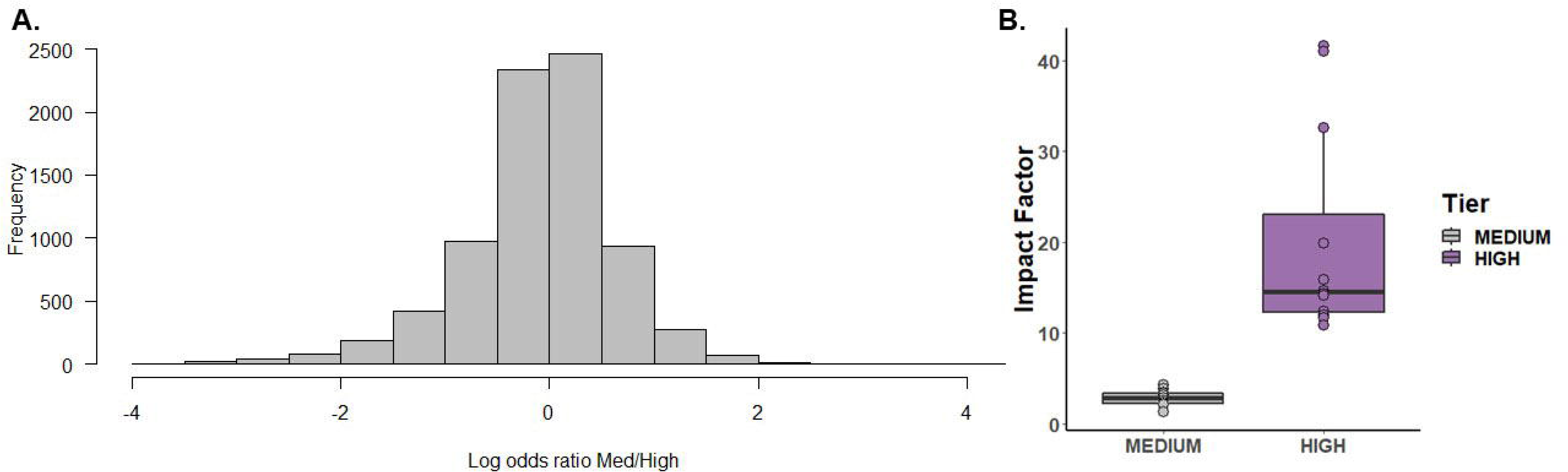
A) The distribution of impact factors among the 12 high- and 13 medium-impact journals selected for comparison. The two outlying high points among the high-impact journals were *Nature* and *Science*. B) The histogram of terms’ log odds ratios. A negative log odds ratio indicates that a term was used more frequently by medium-impact journals.

**Figure 2.**
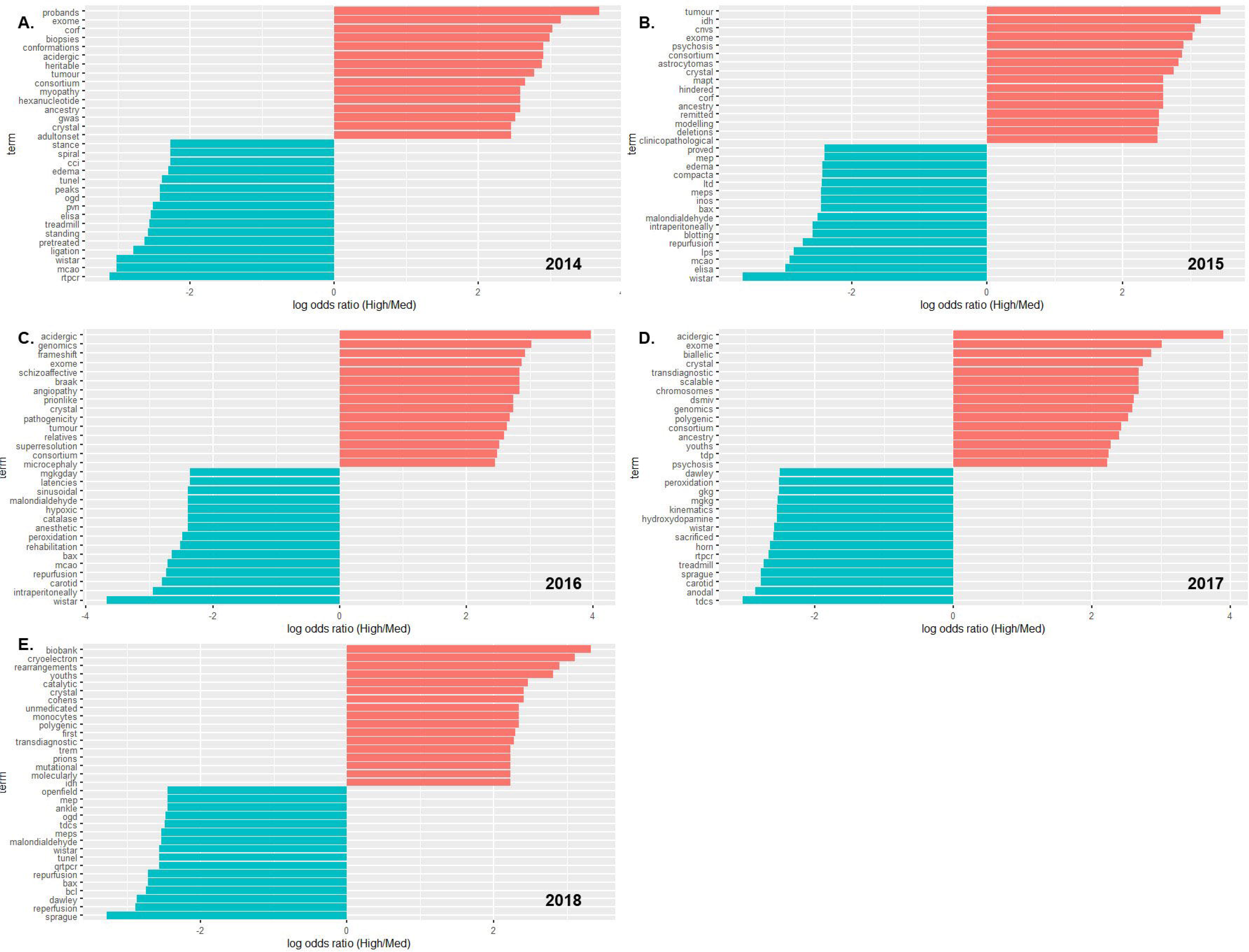
The 15-most biased terms for high- and medium-impact journals on a yearly basis, from 2014 (A) to 2018 (E). Common English stop words and common terms of research / publication were removed. In order to be included, a term had to be used at least 10 times.

**Figure 3.**
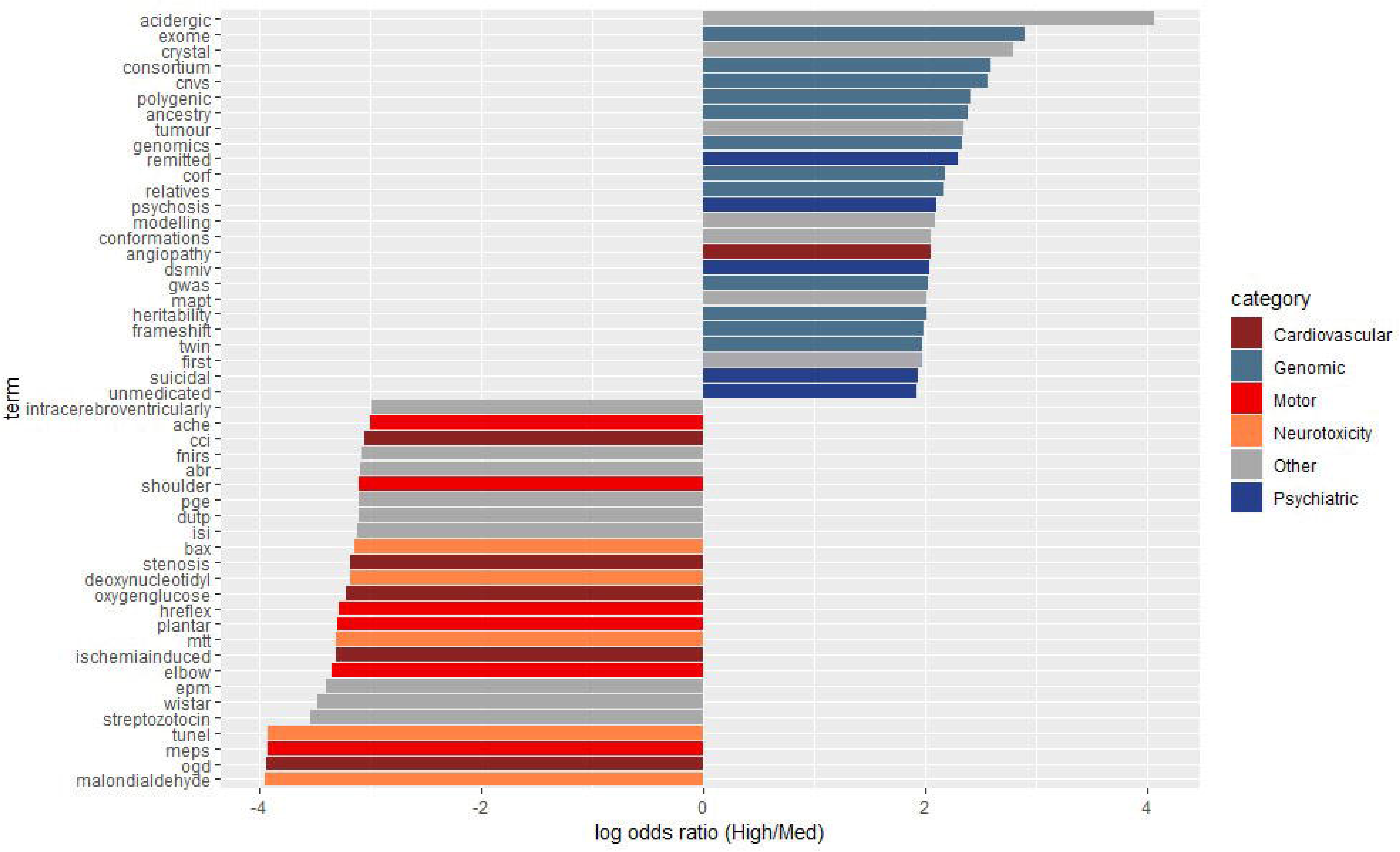
The 25-most biased terms for high- and medium-impact journals across the entire study period from 2014-2018. Terms were then categorized and color-coded as shown in the legend to the right. Common English stop words and common terms of research / publication were removed. In order to be included, a term had to be used at least 50 times.

**Figure 4.**
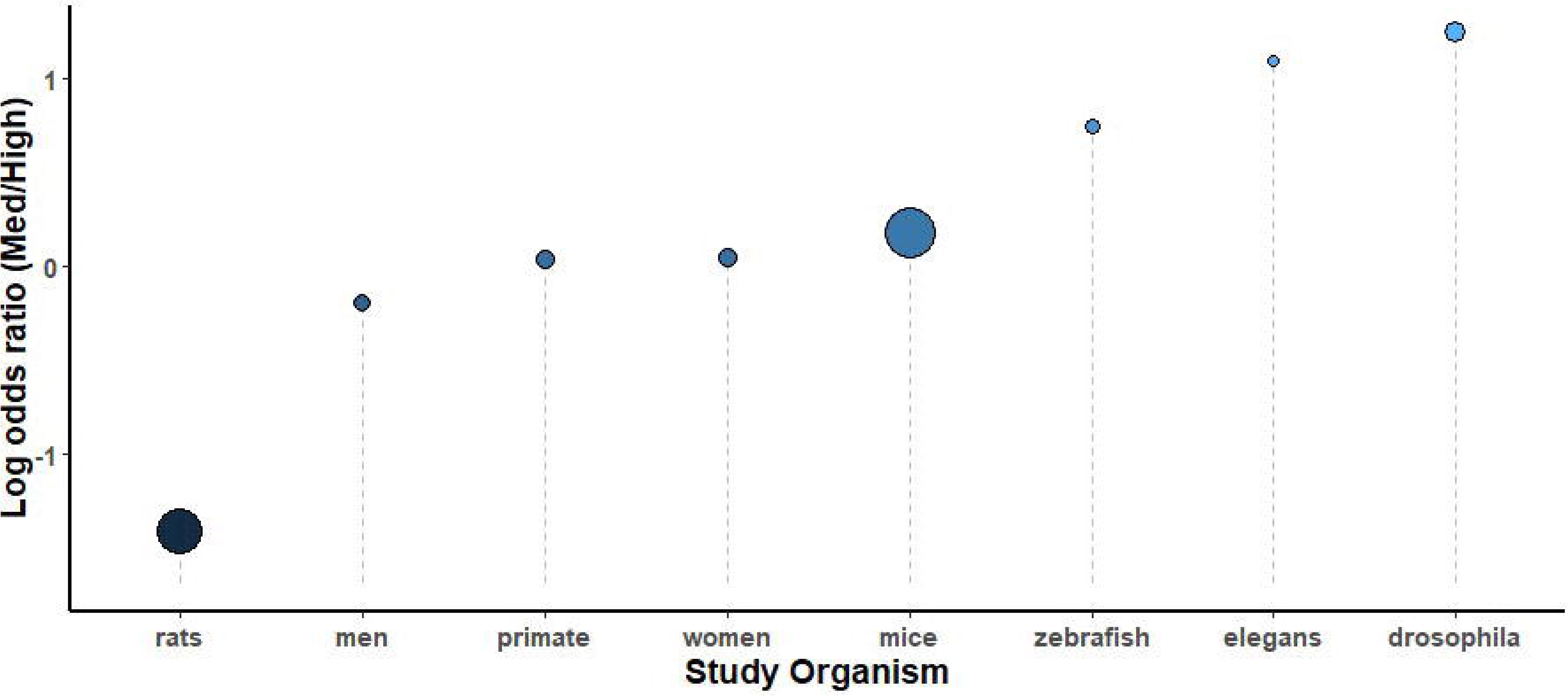
A comparison of study organisms sorted by log odds ratio throughout the study period, 2014-2018. The size of each circle is proportional to the number of instances each term was used, from *C. elegans* (254) to mice (6385).

**Figure 5.**
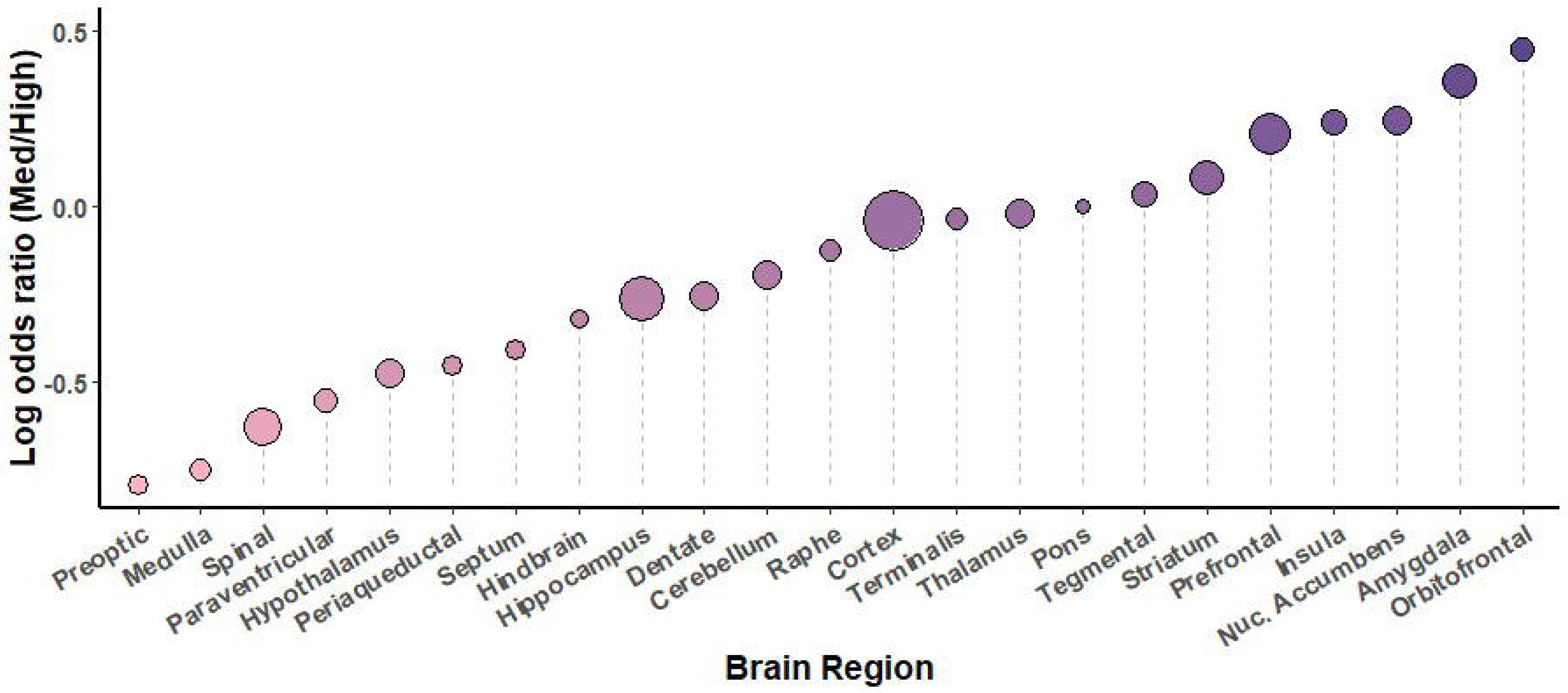
A comparison of brain regions sorted by log odds ratio throughout the study period, 2014-2018. The size of each circle is proportional to the number of instances each term was used, from pons (96) to cortex (9933).

**Figure 6.**
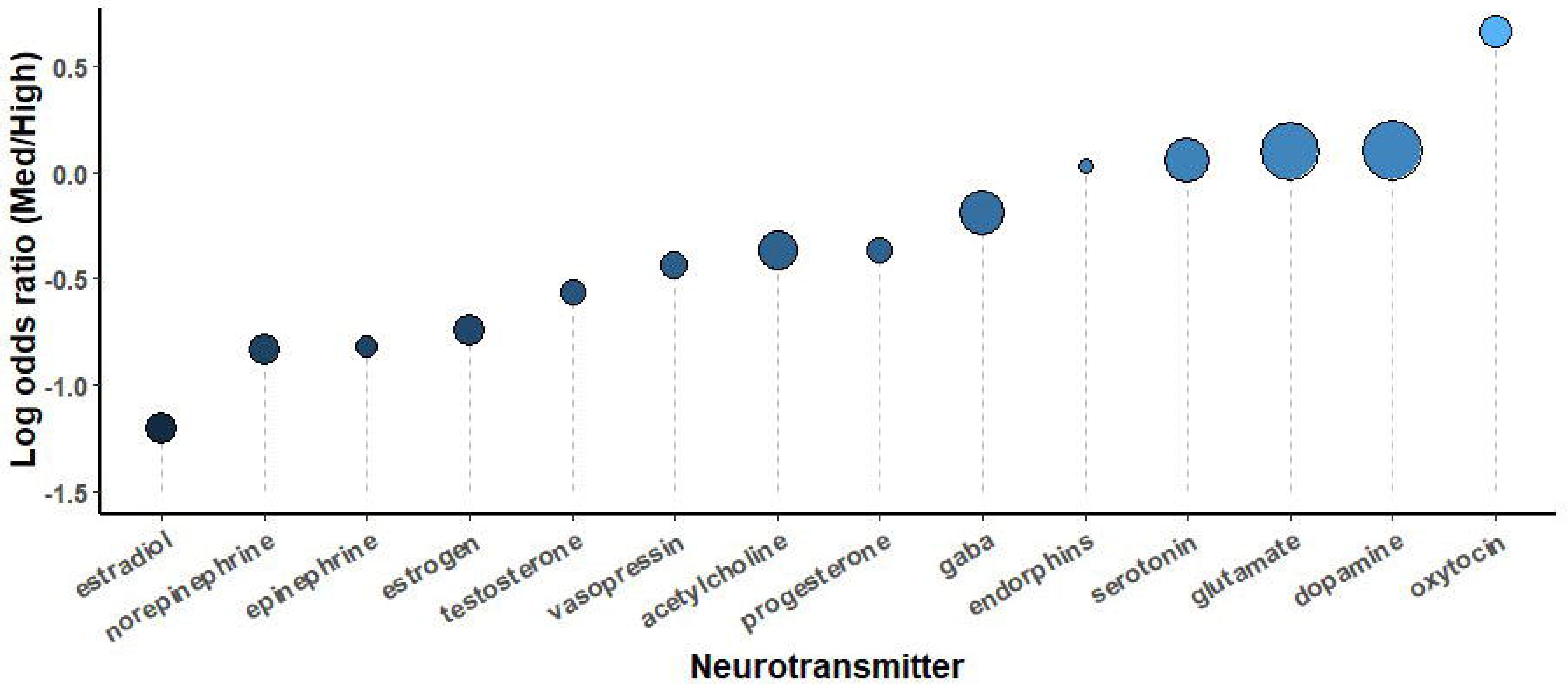
A comparison of neurotransmitters sorted by log odds ratio throughout the study period, 2014-2018. The size of each circle is proportional to the number of instances each term was used, from progesterone (96) to dopamine (2261).

**Figure 7.**
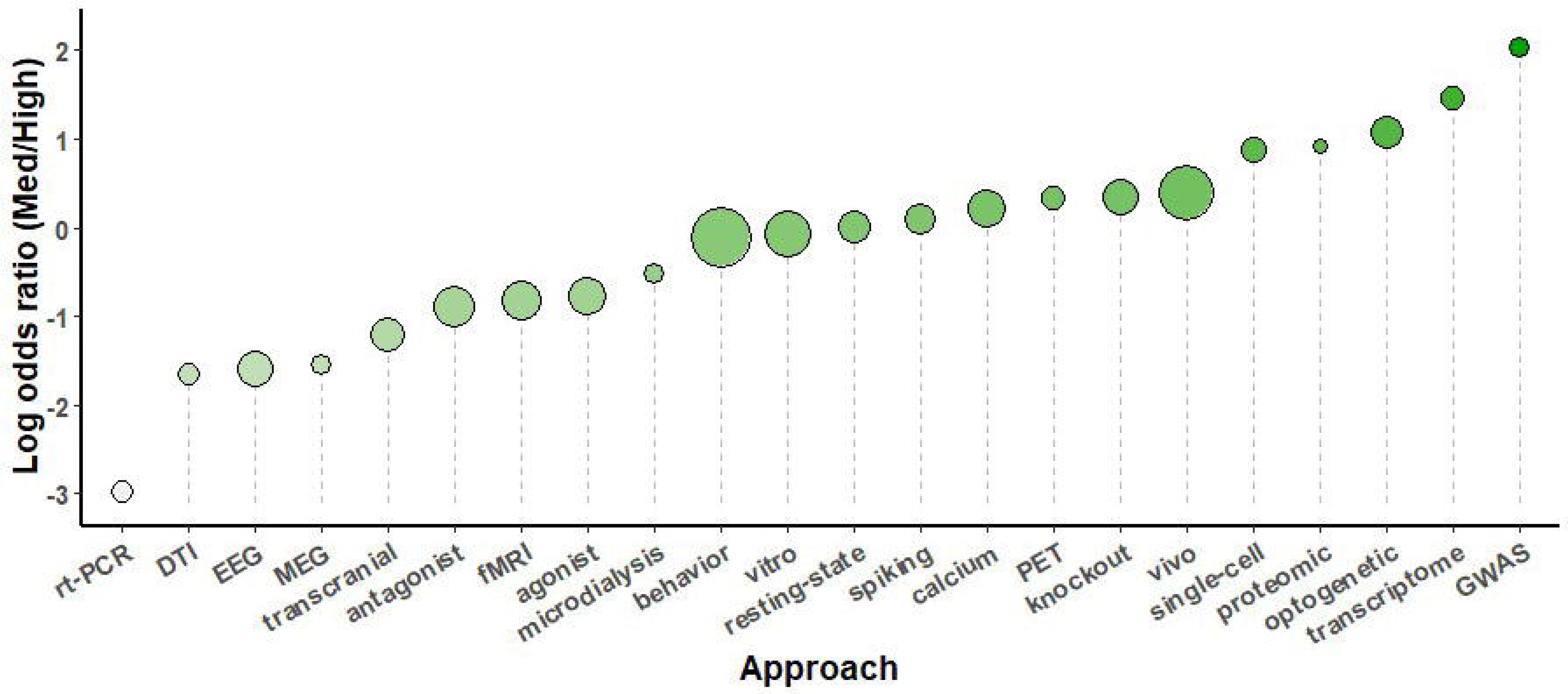
A comparison of methodological approaches sorted by log odds ratio throughout the study period, 2014-2018. The size of each circle is proportional to the number of instances each term was used, from proteomic (132) to behavior (5317).

**Figure 8.**
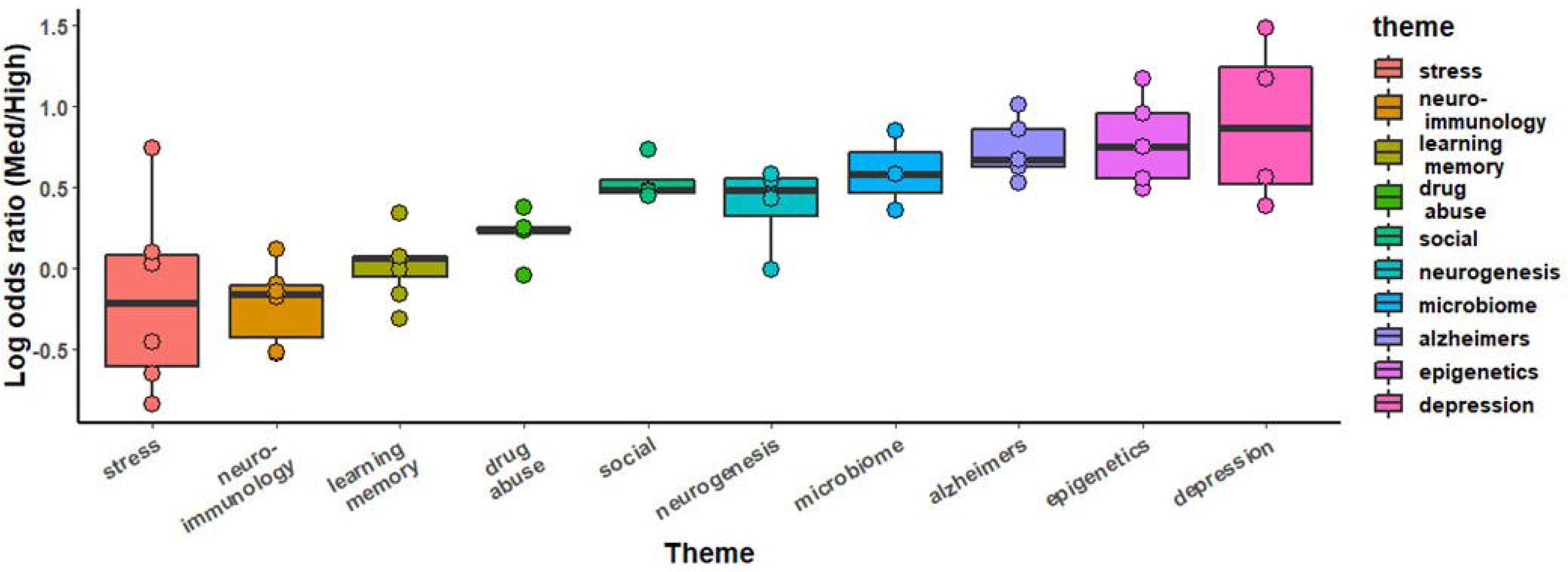
A comparison of broad thematic collections sorted by log odds ratio throughout the study period, 2014-2018. Each theme consists of several related terms, as specified in Table 3.

## DISCUSSION

Several broad themes emerged from the differential use of terms between medium- and high-impact journals. Throughout the study period of 2014 to 2018, there was a clear premium placed on genetic studies, as indicated by the high log odds ratios for terms such as: ‘*ancestry*’, ‘*CNVs*’ (copy number variants), ‘*GWAS*’ (genome-wide association study), ‘*heritable*’, ‘*polygenic*’, and ‘*probands*’. The term ‘*consortium*’ also typically refers to large associations of researchers collaborating on a database of genetic findings. At the same time, terms having to do with psychiatric care also featured prominently in the abstracts of high-impact journals. ‘*DSM-IV’* (Diagnostic and Statistical Manual, 4^th^ edition), ‘*psychosis*’, ‘*suicidal*’, and ‘*un-medicated*’ were all in the top 25 terms with the highest log odds ratio across the study period. These two trends both contributed to the high log odds ratio of ‘*first*’, which occurred in the context of “first episode” and “first-degree relative”. The top most impactful terms were ‘*acidergic*’, ‘*exome*’, and ‘*crystal*’. Although ‘*acidergic*’ was the single most differentially used term among high-impact journals, the more commonly used “gamma-aminobutyric acid”, i.e. GABA, had a log odds ratio of only −0.19 and ‘*gaba-ergic*’, which was used 24x more often, had a log odds ratio of −0.07. ‘*Exome*’ reflects again the preeminence of genetic research within contemporary neuroscience and ‘*crystal*’ refers to x-ray crystallography, a research approach used to determine molecular structure –and one that is notoriously difficult.

The abstracts of medium-impact journals tended to be more specific in terms of their methodology, while high-impact journals tended to use loftier, concept-driven language. The majority of specific neurotransmitters had negative odds ratios, as did many methodological approaches. Terms such as ‘*EMP*’ (elevated plus maze), ‘*MEP*’ (motor-evoked potential), ‘*open-field*’, and ‘*tunel*’ –a marker of apoptosis, all spoke to a more methodologically detailed approach to abstract construction in medium-impact journals. Similarly, details of dosage, such as ‘*g/kg*’, ‘*mg/kg/day*’ and ‘*intraperitoneally*’, were all skewed towards medium-impact journals. It is worth pointing out that these approaches also share the qualities of being widely accessible, relatively inexpensive, long-established, and having low barriers to entry. The ‘*ELISA*’ assay, western ‘*blotting*’, and ‘*rt-PCR*’ are all methods that have not yet been superseded by more advanced approaches, yet each has seen their cache decline as their use has grown more ubiquitous. Similarly, widely-used approaches for human neuroscience showed a consistent bias towards medium-impact journals. ‘*TDCS*’ (transcranial direct current stimulation) and ‘*fNIRS*’ (functional Near-Infrared Spectroscopy) were both consistently among the most medium-impact biased, while DTI, MEG and EEG were all towards the low end of the distribution of log odds ratios compared to other methodologies (Figure 7). This pattern is somewhat in contrast with Yeung et al.’s findings covering 2006-2015 [8], which identified ‘DTI’ and ‘fractional anisotropy’ as high-impact terms (−1.66 and −0.18 log odds ratio in the present work, respectively).

The analysis of research methodologies reinforced the pattern of genetic pre-eminence, with *‘GWAS*’ being the most high-impact biased term. Beyond that was a trio of terms relating to new, challenging and expensive approaches: ‘*transcriptome*’, ‘*optogenetic*’, and ‘*proteomic*’. Both ‘*single-cell*’ and ‘(in) *vivo*’ could refer to a broad range of methods, such as single-cell electrophysiology or in vivo calcium imaging. Indeed, the term ‘*calcium*’ by itself was not far behind. The most frequent methodology term, ‘*behavior*’ was equally used by both medium- and high-impact journals. ‘*CRISPR*’ had a log odds ratio of 2.04 but only occurred 12 times over the study period.

‘*CCI*’ (chronic constrictive injury), ‘*edema*’, ‘*MCAO*’ (middle cerebral artery occlusion), ‘*oxygen-glucose*’, ‘*oxygen-glucose-deprivation*’, ‘*reperfusion*’, and ‘*stenosis*’ were terms biased towards medium-impact journals that all pertain to the cardiovascular pathology of stroke. These terms were spread across several medium-impact journals, which suggests a broad pattern. Indeed, previous analysis of neuroscience articles from 2006-2015 also identified ischemic stroke as having consistently low citation impacts, along with multiple sclerosis and intracerebral hemorrhage [8]. It was especially curious then that ‘*angiopathy*’ shows up in the list of most high-impact biased terms. A close examination of abstracts containing ‘*angiopathy*’ revealed that they tended to be focused on Alzheimer’s pathology.

Another trend among the medium-impact journal abstracts was the high frequency of rat strain names. ‘*Wistar*’ and ‘*Sprague-Dawley*’ consistently featured in the top 15 most differentiated terms in favor of medium-impact journals. Among study organisms, the most frequent terms were rats and mice, although the phrasing of abstracts with regards to human subjects can be quite broad, and thus we are left without a direct comparison of the percentage of usage of each organism. The overall pattern that emerges is that of a U-shaped curve if impact were the y-axis and a continuum of translatability were the x-axis. Studies of human psychiatric subjects were biased towards high-impact journals, as were studies of zebrafish, *C. elegans* and drosophila. Mammalian models, particularly rats, are left in the trough of this curve. To wit, within the “stress” theme, ‘*cortisol*’ (the human glucocorticoid) had a higher log odds ratio than ‘*corticosterone*’ (the rodent analogue), 0.74 vs. −0.68, respectively.

Among neurotransmitters, most terms were biased towards medium-impact journals, with only ‘*oxytocin*’ showing a substantially positive log odds ratio. On the other hand, themes were mostly biased towards high-impact journals. The pattern among themes remained similar when the threshold for term inclusion was dropped from 10 instances / year to 10 instances across the entire 5-year span, with the notable exception of the Microbiome theme. Within Microbiome, the lowered threshold led to the inclusion of ‘microbiome’, ‘germ-free’, and ‘microbial’, which each had log odds ratios greater than 1.46. Among brain regions, there was a generally caudal-rostral progression in terms of log odds ratio, with several exceptions. As expected, ‘*cortex*’ was widely used, ‘*hippocampus*’ was the most commonly used specific region, and ‘*orbitofrontal*’ (cortex), a region almost exclusively studied in humans, the most high-impact biased.

Previous work by Yeung et al. identified ‘autism’, ‘meta-analysis’, ‘functional connectivity’, ‘default mode network’ and ‘neuroimaging’ as the most consistent terms associated with garnering the most citations from 2006 to 2015 [8]. From 2012 to 2015, the terms ‘melatonin’, ‘microglia’, and ‘neurofibrillary tangle’ each also emerged as high impact [8]. Although direct comparison to present findings is limited, as the algorithm used here considered only single words, these terms’ performance from 2014 to 2018 did not distinguish them as exceptionally high-impact associated: ‘*autism*’ (1.06), ‘*melatonin*’ (−1.72), ‘*meta-analysis*’ (1.0), ‘*microglia*’ (0.08), ‘*neurofibrillary*’ (0.89), and ‘*neuroimaging*’ (0.58).

Such a gross overview of an entire field is sure to come with caveats. First, it should be said that the secondary analyses were unavoidably influenced by the author’s interests and presumably by his biases as well. A more molecular author may have chosen to compare cell organelles rather than brain regions. A more statistically-minded author would have examined language relating to techniques of analysis. These analyses could also have been limited by the selection of journals. Although every effort was made to ensure this selection was representative, there could still be lurking biases within their composition. The very fact that some of neuroscience’s highest-impact journals consist entirely of review articles is likely to have influenced the text of their abstracts. This is likely to have contributed to the observation that high-impact journals were more likely to use concept-based language and to focus less on methods-based language. Similarly, two of the nine high-impact journals had “psychiatry” in the title (*Biological Psychiatry* and *Molecular Psychiatry*), so it should not come as a great surprise that terms related to psychiatry should feature highly in the list of high-impact biased. A more rigorous set of analyses could be done if impact factor was considered as a continuous variable rather than being collapsed into two categories as done here. Finally, it should also be noted that the citation impact of an individual researcher’s early work is not a reliable predictor of career persistence [9].

The field of neuroscience has long ago outgrown informal assessments and conventional wisdom. Only by monitoring the content of publications can we maintain an objective perspective on the field’s priorities. Given the discrepancies between the present findings and the few comparable analyses of the past, it would appear that was in vogue one year may be superseded quickly. That this churn in the field’s interest can occur within the span of a graduate student’s training period should have major implications for career mentorship. Analyses like those presented here should be carried our regularly, and ideally, in the future with more rigorous, completeness, and sophistication.

## Supporting information

Abstract Data

Analysis Code

