## Supplementary material for "Corpus Colossal: A Bibliometric Analysis of Neuroscience Abstracts and Impact Factor": Analysis Code: Impact Graph.R

```
JournalHomeGrid <- read_excel("JournalHomeGrid.xlsx", skip = 2)
JournalHomeGrid <- JournalHomeGrid[which(!is.na(JournalHomeGrid$Tier)),]
JournalHomeGrid$`Journal Impact Factor` <-as.numeric(JournalHomeGrid$`Journal Impact Factor`)
JournalHomeGrid$Tier <- factor(JournalHomeGrid$Tier, levels = c('MEDIUM','HIGH'))


e <- ggplot(JournalHomeGrid, aes(x = Tier, y = `Journal Impact Factor`)) + ylab("Impact Factor") + xlab("") + theme_classic()
e <- e + geom_boxplot(aes(fill = Tier), size = 1)  
e <- e + scale_fill_manual(values = c("#BFBFBF", "#9C71AB"))
# and now the dots
e <- e + geom_point(aes(fill=Tier), colour="black",shape=21,size=4,show.legend=FALSE, position=position_dodge(.75))
e <- e + scale_color_manual(values = c("#BFBFBF", "#9C71AB"))
# set up the text:
e <- e + theme(text = element_text(size=20, face="bold"), axis.line = element_line(color="black", size=1, linetype="solid"))
```
