## Supplementary material for "Corpus Colossal: A Bibliometric Analysis of Neuroscience Abstracts and Impact Factor": Analysis Code: Master Analysis.R

```
# CORF = C9ORF72
# MEP = motor evoked potential
# OGD = oxygen glucose deprivation
# MWM = Morris water maze
# MTT = cell metabolic activity assay
# BAX = apoptosis signal
# FNIRS = Functional Near Infrared Spectroscopy
# CCI = chronic constrictive injury
# MAPT = gene encoding tau protein relevant to Alzheimer's
# ABR = auditory brain stem responses 
# PGE = prostaglandin E2
# dUTP = deoxyuridine triphosphate
# MCAO = middle cerebral artery occlusion
# Malondialdehyde = result of lipid peroxidation
# ISI = inter-stimulus interval / inter-spike interval

# MDA should be NMDA


rm(list = ls())
setwd("D:/Box Sync/Research/Meta Review")

load("D:/Box Sync/Research/Meta Review/2018 Data.RData")
masterhigh <- highabstracts
masterlow <- lowabstracts
load("D:/Box Sync/Research/Meta Review/2017 Data.RData")
masterhigh <- c(masterhigh,highabstracts)
masterlow <- c(masterlow,lowabstracts)
load("D:/Box Sync/Research/Meta Review/2016 Data.RData")
masterhigh <- c(masterhigh,highabstracts)
masterlow <- c(masterlow,lowabstracts)
load("D:/Box Sync/Research/Meta Review/2015 Data.RData")
masterhigh <- c(masterhigh,highabstracts)
masterlow <- c(masterlow,lowabstracts)
load("D:/Box Sync/Research/Meta Review/2014 Data.RData")
masterhigh <- c(masterhigh,highabstracts)
masterlow <- c(masterlow,lowabstracts)

scienceywords = c("may","study","group","effect","effects","increased","decreased","significant")
scienceywords = c(scienceywords, "significantly","higher","lower","compared","control","including","use")
scienceywords = c(scienceywords, "administration","response","groups","different","present","data","one")
scienceywords = c(scienceywords, "two","whether","day","present","reduced","evidence","first","role","right")
scienceywords = c(scienceywords, "elsevier","found","involved","used","suggest","analysis","can","different")
scienceywords = c(scienceywords, "treatment", "associated", "increase", "time", "hours", "differences","activity")
scienceywords = c(scienceywords, "factors", "changes", "respectively", "rights", "trials", "function","mechanisms")
scienceywords = c(scienceywords, "showed", "findings", "min", "days", "well", "received", "however", "also")
scienceywords = c(scienceywords, "observed", "studies", "results", "using", "also", "less", "without", "rate")
scienceywords = c(scienceywords, "high", "level", "research", "measured", "reserved", "patients", "within")
scienceywords = c(scienceywords, "statement", "wiley", "ibro", "sons", "federation", "noteworthy", "periodicals")
scienceywords = c(scienceywords, "john", "societies", "inâ", "isâ", "andâ", "toâ", "abstractcopyright", "etâ", "skill")
scienceywords = c(scienceywords, "extension", "japan", "ireland", "instructed", "presumably", "prepared", "chapter")
scienceywords = c(scienceywords, "reservedthis", "inc", "theâ", "colleagues","incâ", "maximal", "meanwhile", "alternating")
scienceywords = c(scienceywords, "journal", "authors", "ofâ", "comp", "neurol", "american", "imply", "exp", "european")
scienceywords = c(scienceywords, "righthanded", "biology", "press", "commentary", "author", "university", "please")
scienceywords = c(scienceywords, "society", "permissions", "journalspermissionsoupcom", "email", "oxford", "guarantors")
scienceywords = c(scienceywords, "behalf", "examples", "biological", "firestone", "doiaww", "challenges", "inclusions")
scienceywords = c(scienceywords, "morsella", "gane", "experiment","pathwaycopyright","ilî","doiawx","leibovich","institute")
scienceywords = c(scienceywords, "kalisch", "duarte", "lane", "doibrainawy","mahr", "csibra", "psychiatry","advancement")
scienceywords = c(scienceywords, "australian", "ackermann", "shanks", "newell", "bentley", "pepper", "nettle","sciencey")
scienceywords = c(scienceywords, "christiansen", "cop", "copyright", "lake","kline","klines","sci","sweden","over")
scienceywords = c(scienceywords, "entitled","doibrainaww","diseaseâ","science","neurosci","doi","licensee","claim")
scienceywords = c(scienceywords, "government","exclusive","neurophysiol")

breakwords <- function(corpus){
  corpus <- tm_map(corpus, content_transformer(gsub), pattern = "spraguedawley", replacement = "sprague dawley")
  corpus <- tm_map(corpus, content_transformer(gsub), pattern = "attentiondeficithyperactivity", replacement = "attention deficit hyperactivity")
  corpus <- tm_map(corpus, content_transformer(gsub), pattern = "singlenucleotide", replacement = "single nucleotide")
  corpus <- tm_map(corpus, content_transformer(gsub), pattern = "machinelearning", replacement = "machine learning")
  corpus <- tm_map(corpus, content_transformer(gsub), pattern = "genomewide", replacement = "genome wide")
  corpus <- tm_map(corpus, content_transformer(gsub), pattern = "firstepisode", replacement = "first episode")
  corpus <- tm_map(corpus, content_transformer(gsub), pattern = "firstdegree", replacement = "first degree")
  corpus <- tm_map(corpus, content_transformer(gsub), pattern = "obsessivecompulsive", replacement = "obsessive compulsive")
  corpus <- tm_map(corpus, content_transformer(gsub), pattern = "lossoffunction", replacement = "loss of function")
  corpus <- tm_map(corpus, content_transformer(gsub), pattern = "placebocontrolled", replacement = "placebo controlled")
  corpus <- tm_map(corpus, content_transformer(gsub), pattern = "tnfî", replacement = "tnf")
  corpus <- tm_map(corpus, content_transformer(gsub), pattern = "ratscopyright", replacement = "rats copyright")
  corpus <- tm_map(corpus, content_transformer(gsub), pattern = "ischemiareperfusion", replacement = "ischemia repurfusion")
  corpus <- tm_map(corpus, content_transformer(gsub), pattern = "wholeexome", replacement = "whole exome")
  corpus <- tm_map(corpus, content_transformer(gsub), pattern = "patientderived", replacement = "patient derived")
  corpus <- tm_map(corpus, content_transformer(gsub), pattern = "pairedpulse", replacement = "paired pulse")
  corpus <- tm_map(corpus, content_transformer(gsub), pattern = "nearinfrared", replacement = "near infrared")
  corpus <- tm_map(corpus, content_transformer(gsub), pattern = "atrisk", replacement = "at risk")
  corpus <- tm_map(corpus, content_transformer(gsub), pattern = "wholegenome", replacement = "whole genome")
  corpus <- tm_map(corpus, content_transformer(gsub), pattern = "diseaseassociated", replacement = "disease associated")
  corpus <- tm_map(corpus, content_transformer(gsub), pattern = "schizophreniacopyright", replacement = "schizophrenia copyright")
  corpus <- tm_map(corpus, content_transformer(gsub), pattern = "populationbased", replacement = "population based")
  corpus <- tm_map(corpus, content_transformer(gsub), pattern = "celladhesion", replacement = "cell adhesion")
  corpus <- tm_map(corpus, content_transformer(gsub), pattern = "asdcopyright", replacement = "asd copyright")
  corpus <- tm_map(corpus, content_transformer(gsub), pattern = "ligandbinding", replacement = "ligand binding")
  corpus <- tm_map(corpus, content_transformer(gsub), pattern = "overrepresented", replacement = "over represented")
  corpus <- tm_map(corpus, content_transformer(gsub), pattern = "oestrogen", replacement = "estrogen")  
  corpus <- tm_map(corpus, content_transformer(gsub), pattern = "ageing", replacement = "aging")
  corpus <- tm_map(corpus, content_transformer(gsub), pattern = "analog", replacement = "analogue")
  corpus <- tm_map(corpus, content_transformer(gsub), pattern = "tumours", replacement = "tumour")
  corpus <- tm_map(corpus, content_transformer(gsub), pattern = "centres", replacement = "centers")
  corpus <- tm_map(corpus, content_transformer(gsub), pattern = "neighbouring", replacement = "neighboring")
  corpus <- tm_map(corpus, content_transformer(gsub), pattern = "dosedependently", replacement = "dose dependently")
  corpus <- tm_map(corpus, content_transformer(gsub), pattern = "shamoperated", replacement = "sham operated")
  corpus <- tm_map(corpus, content_transformer(gsub), pattern = "repurfusion", replacement = "reperfusion")
  corpus <- tm_map(corpus, content_transformer(gsub), pattern = "meps", replacement = "mep")
  
  #corpus <- tm_map(corpus, content_transformer(gsub), pattern = "spraguedawley", replacement = "sprague")
  return(corpus)
}

abcorp <- Corpus(VectorSource(masterhigh))

abmap <- tm_map(abcorp, PlainTextDocument)
abmap <- tm_map(abmap, removePunctuation)
abmap <- tm_map(abmap, removeWords, stopwords('english'))
#abmap <- tm_map(abmap, stemDocument)
# I personally don't like to stem the words

termy <- clean_corpus(abmap)
termy <- breakwords(termy)
term_dtm <- DocumentTermMatrix(termy)
# According to https://www.tidytextmining.com/dtm.html 'Converting to and from...' this is the step
# where we need to convert from corpus into text data using the 'tidy' function

high_words <- tidy(term_dtm)
high_words$tier <- 'High'
high_words$total <- sum(high_words$count)

abcorp <- Corpus(VectorSource(masterlow))

abmap <- tm_map(abcorp, PlainTextDocument)
abmap <- tm_map(abmap, removePunctuation)
abmap <- tm_map(abmap, removeWords, stopwords('english'))

termy <- clean_corpus(abmap)
termy <- breakwords(termy)
term_dtm <- DocumentTermMatrix(termy)

low_words <- tidy(term_dtm)
low_words$tier <- 'Low'
low_words$total <- sum(low_words$count)

abstracts <- bind_rows(low_words, high_words)

frequency <- abstracts %>% 
  group_by(tier) %>% 
  count(term, sort = TRUE) %>% 
  left_join(abstracts %>% 
              group_by(tier) %>% 
              summarise(total = n())) %>%
  mutate(freq = n/total)

frequency <- frequency %>% 
  select(tier, term, freq) %>% 
  spread(tier, freq) %>%
  arrange(Low, High)

# note that we increase the cutoff from 10 to 50 because this is the master list
word_ratios <- abstracts %>%
  count(term, tier) %>%
  group_by(term) %>%
  filter(sum(n) >= 50) %>%
  filter(!str_detect(term, "copyright")) %>%
  ungroup() %>%
  spread(tier, n, fill = 0) %>%
  mutate(instance = as.character(High + Low)) %>%
  mutate_if(is.numeric, funs((. + 1) / (sum(.) + 1))) %>%
  mutate(logratio = log(High / Low)) %>%
  arrange(desc(logratio))
word_ratios$instance <- as.numeric(word_ratios$instance)

word_ratios %>%
  group_by(logratio < 0) %>%
  top_n(25, abs(logratio)) %>%
  ungroup() %>%
  mutate(term = reorder(term, logratio)) %>%
  ggplot(aes(term, logratio, fill = logratio < 0)) +
  geom_col(show.legend = FALSE) +
  coord_flip() +
  ylab("log odds ratio (High/Med)") +
  scale_fill_discrete(name = "", labels = c("High", "Med"))


# to view and weed out nonsense words, check dump then add them to sciencywords
dump <- word_ratios %>% group_by(logratio < 0) %>% top_n(50, abs(logratio))

# now let's assign each term a category
codedwords <- word_ratios
codedwords %>%
  group_by(logratio < 0) %>%
  top_n(25, abs(logratio)) %>%
  ungroup() %>%
  mutate(term = reorder(term, logratio))
# add categories to the terms
codedwords <- read.csv("Top Terms.csv")
codedwords %>%
  group_by(logratio < 0) %>%
  top_n(25, abs(logratio)) %>%
  ungroup() %>%
  mutate(term = reorder(term, logratio)) %>%
  ggplot(aes(term, logratio, fill = category)) +
  geom_col() +
  scale_fill_manual(values=c("brown4", "skyblue4", "red2",  "sienna1", "dark gray", "royalblue4")) +
  coord_flip() +
  ylab("log odds ratio (High/Med)")

# to look up the odds ratio of a specific term
# word_ratios[match('optogenetics',word_ratios$term),]

nam <- 'Master Data.RData'
save.image(file = nam)

hist(word_ratios$logratio, main="", xlab="Log odds ratio Med/High ", col="gray", las=1,breaks=20)

###
###
###

#let's find some of these weird words and see what they mean...
for (i in 1:length(highabstracts)){
  
  abcorp <- Corpus(VectorSource(highabstracts[i]))
  
  abmap <- tm_map(abcorp, PlainTextDocument)
  abmap <- tm_map(abmap, removePunctuation)
  abmap <- tm_map(abmap, removeWords, stopwords('english'))
  
  termy <- clean_corpus(abmap)
  termy <- breakwords(termy)
  
  if(length(grep("exclusive", termy))>0)
    print(highabstracts[i])
}

for (i in 1:length(lowabstracts)){
  
  abcorp <- Corpus(VectorSource(lowabstracts[i]))
  
  abmap <- tm_map(abcorp, PlainTextDocument)
  abmap <- tm_map(abmap, removePunctuation)
  abmap <- tm_map(abmap, removeWords, stopwords('english'))
  
  termy <- clean_corpus(abmap)
  termy <- breakwords(termy)
  
  if(length(grep("malondialdehyde",termy))>0)
    print(lowabstracts[i])
}

###
###
###
```
