## Supplementary material for "Corpus Colossal: A Bibliometric Analysis of Neuroscience Abstracts and Impact Factor": Analysis Code: Review Abstract Miner ALL YEARS.R

```
rm(list = ls())

yr <- 2014
setwd("D:/Box Sync/Research/Meta Review")

highs <- c('Nature Reviews Neuroscience','Nature Neuroscience','Annual Review of Neuroscience','Neuron')
highs <- c(highs,'Molecular Psychiatry','Biological Psychiatry', 'Acta Neuropathologica', 'Progress in Neurobiology', 'Brain')
# 'Trends in Neurosciences' has too short of abstracts

potentials <- c('Nature', 'Science', 'Nature Communications')

keywords = c("brain","brainstem","neuron","neurons","neural","synapse","synaptic","neuroscience","neurotransmitter")
keywords = c(keywords,"amygdala","axonal","dendrite","axons","dendrites","hippocampus","hypothalamus","thalamus")
keywords = c(keywords, "myelination","myelin","cerebellum","sulcus","gyrus")
keywords = c(keywords,"dopamine" , "serotonin" , "gaba" , "glutamate" , "acetylcholine" , "endorphins" , "oxytocin" , "vasopressin")
keywords <- sort(keywords)

lows <- c('Journal of Neurophysiology','Neuroscience','Frontiers in Neuroscience','Brain Structure and Function')
lows <- c(lows,'Brain Research','Neuroscience Bulletin','Brain and Behavior','NeuroReport','European Journal of Neuroscience')
lows <- c(lows,'Brain Research Bulletin','BMC Neuroscience','Neuroscience Research','Neuroscience Letters')

scienceywords = c("may","study","group","effect","effects","increased","decreased","significant")
scienceywords = c(scienceywords, "significantly","higher","lower","compared","control","including","use")
scienceywords = c(scienceywords, "administration","response","groups","different","present","data","one")
scienceywords = c(scienceywords, "two","whether","day","present","reduced","evidence","first","role","right")
scienceywords = c(scienceywords, "elsevier","found","involved","used","suggest","analysis","can","different")
scienceywords = c(scienceywords, "treatment", "associated", "increase", "time", "hours", "differences","activity")
scienceywords = c(scienceywords, "factors", "changes", "respectively", "rights", "trials", "function","mechanisms")
scienceywords = c(scienceywords, "showed", "findings", "min", "days", "well", "received", "however", "also")
scienceywords = c(scienceywords, "observed", "studies", "results", "using", "also", "less", "without", "rate")
scienceywords = c(scienceywords, "high", "level", "research", "measured", "reserved", "patients", "within")
scienceywords = c(scienceywords, "statement", "wiley", "ibro", "sons", "federation", "noteworthy", "periodicals")
scienceywords = c(scienceywords, "john", "societies", "inâ", "isâ", "andâ", "toâ", "abstractcopyright", "etâ", "skill")
scienceywords = c(scienceywords, "extension", "japan", "ireland", "instructed", "presumably", "prepared", "chapter")
scienceywords = c(scienceywords, "reservedthis", "inc", "theâ", "colleagues","incâ", "maximal", "meanwhile", "alternating")
scienceywords = c(scienceywords, "journal", "authors", "ofâ", "comp", "neurol", "american", "imply", "exp", "european")
scienceywords = c(scienceywords, "righthanded", "biology", "press", "commentary", "author", "university", "please")
scienceywords = c(scienceywords, "society", "permissions", "journalspermissionsoupcom", "email", "oxford", "guarantors")
scienceywords = c(scienceywords, "behalf", "examples", "biological", "firestone", "doiaww", "challenges", "inclusions")
scienceywords = c(scienceywords, "morsella", "gane", "experiment","pathwaycopyright","ilî","doiawx","leibovich","institute")
scienceywords = c(scienceywords, "kalisch", "duarte", "lane", "doibrainawy","mahr", "csibra", "psychiatry","advancement")
scienceywords = c(scienceywords, "australian", "ackermann", "shanks", "newell", "bentley", "pepper", "nettle","sciencey")
scienceywords = c(scienceywords, "christiansen", "cop", "copyright", "lake","kline","klines","sci","sweden","over")
scienceywords = c(scienceywords, "entitled","doibrainaww","diseaseâ","science","neurosci","doi","licensee","claim")
scienceywords = c(scienceywords, "government")

addpotents <- function(highabstracts)
{
  potentids <- getPMIDsByKeyWords(journal = potentials[1],dFrom = yr,
                                  dTo = yr, n = 10000, https = TRUE )
  for (i in 2:length(potentials)) {
    potentids <- c( potentids, getPMIDsByKeyWords( journal = potentials[i],dFrom = yr,dTo = yr,
                                                   n = 10000, https = TRUE ))
    Sys.sleep(0.5)
  }
  
  for (j in 1:length(potentids)) {
    Sys.sleep(0.25)
    beep_on_error(tryCatch({
      potabstract <- getAbstracts(potentids[j])
    }, error = function(e) {
      Sys.sleep(0.75)
      potabstract <- getAbstracts(potentids[j])
    }), sound = 1)
    
    if (length(potabstract) > 0) {
      if (sum(sapply(keywords, grepl, potabstract)) > 0) {
        highabstracts <- c(highabstracts, potabstract)
      }
    }
    print(j)
    print(Sys.time())
  }
  return(highabstracts)
}

clean_corpus <- function(corpus){
  corpus <- tm_map(corpus, stripWhitespace)
  corpus <- tm_map(corpus, removeNumbers)
  corpus <- tm_map(corpus, removePunctuation)
  corpus <- tm_map(corpus, content_transformer(tolower))
  corpus <- tm_map(corpus, removeWords, c(stopwords("english"),scienceywords))
  return(corpus)
}

breakwords <- function(corpus){
  corpus <- tm_map(corpus, content_transformer(gsub), pattern = "spraguedawley", replacement = "sprague dawley")
  corpus <- tm_map(corpus, content_transformer(gsub), pattern = "attentiondeficithyperactivity", replacement = "attention deficit hyperactivity")
  corpus <- tm_map(corpus, content_transformer(gsub), pattern = "singlenucleotide", replacement = "single nucleotide")
  corpus <- tm_map(corpus, content_transformer(gsub), pattern = "machinelearning", replacement = "machine learning")
  corpus <- tm_map(corpus, content_transformer(gsub), pattern = "genomewide", replacement = "genome wide")
  corpus <- tm_map(corpus, content_transformer(gsub), pattern = "firstepisode", replacement = "first episode")
  corpus <- tm_map(corpus, content_transformer(gsub), pattern = "firstdegree", replacement = "first degree")
  corpus <- tm_map(corpus, content_transformer(gsub), pattern = "obsessivecompulsive", replacement = "obsessive compulsive")
  corpus <- tm_map(corpus, content_transformer(gsub), pattern = "lossoffunction", replacement = "loss of function")
  corpus <- tm_map(corpus, content_transformer(gsub), pattern = "placebocontrolled", replacement = "placebo controlled")
  corpus <- tm_map(corpus, content_transformer(gsub), pattern = "tnfî", replacement = "tnf")
  corpus <- tm_map(corpus, content_transformer(gsub), pattern = "ratscopyright", replacement = "rats copyright")
  corpus <- tm_map(corpus, content_transformer(gsub), pattern = "ischemiareperfusion", replacement = "ischemia repurfusion")
  corpus <- tm_map(corpus, content_transformer(gsub), pattern = "wholeexome", replacement = "whole exome")
  corpus <- tm_map(corpus, content_transformer(gsub), pattern = "patientderived", replacement = "patient derived")
  corpus <- tm_map(corpus, content_transformer(gsub), pattern = "pairedpulse", replacement = "paired pulse")
  corpus <- tm_map(corpus, content_transformer(gsub), pattern = "nearinfrared", replacement = "near infrared")
  corpus <- tm_map(corpus, content_transformer(gsub), pattern = "atrisk", replacement = "at risk")
  corpus <- tm_map(corpus, content_transformer(gsub), pattern = "wholegenome", replacement = "whole genome")
  corpus <- tm_map(corpus, content_transformer(gsub), pattern = "diseaseassociated", replacement = "disease associated")
  corpus <- tm_map(corpus, content_transformer(gsub), pattern = "schizophreniacopyright", replacement = "schizophrenia copyright")
  corpus <- tm_map(corpus, content_transformer(gsub), pattern = "populationbased", replacement = "population based")
  corpus <- tm_map(corpus, content_transformer(gsub), pattern = "celladhesion", replacement = "cell adhesion")
  corpus <- tm_map(corpus, content_transformer(gsub), pattern = "asdcopyright", replacement = "asd copyright")
  corpus <- tm_map(corpus, content_transformer(gsub), pattern = "ligandbinding", replacement = "ligand binding")
  corpus <- tm_map(corpus, content_transformer(gsub), pattern = "overrepresented", replacement = "over represented")
  corpus <- tm_map(corpus, content_transformer(gsub), pattern = "oestrogen", replacement = "estrogen")  
  corpus <- tm_map(corpus, content_transformer(gsub), pattern = "ageing", replacement = "aging")
  corpus <- tm_map(corpus, content_transformer(gsub), pattern = "analog", replacement = "analogue")
  corpus <- tm_map(corpus, content_transformer(gsub), pattern = "tumours", replacement = "tumour")
  corpus <- tm_map(corpus, content_transformer(gsub), pattern = "centres", replacement = "centers")
  corpus <- tm_map(corpus, content_transformer(gsub), pattern = "neighbouring", replacement = "neighboring")
  corpus <- tm_map(corpus, content_transformer(gsub), pattern = "dosedependently", replacement = "dose dependently")
  corpus <- tm_map(corpus, content_transformer(gsub), pattern = "shamoperated", replacement = "sham operated")
  corpus <- tm_map(corpus, content_transformer(gsub), pattern = "repurfusion", replacement = "reperfusion")
  
  #corpus <- tm_map(corpus, content_transformer(gsub), pattern = "spraguedawley", replacement = "sprague")
  return(corpus)
}

highids <- getPMIDsByKeyWords(journal = highs[1], dFrom = yr, dTo = yr, n = 10000, https = TRUE)
for(i in 2:length(highs)){
  highids <- c(highids,getPMIDsByKeyWords(journal = highs[i], dFrom = yr, dTo = yr, n = 10000, https = TRUE))
  Sys.sleep(0.5)
}
potentids <- getPMIDsByKeyWords(journal = potentials[1], dFrom = yr, dTo = yr, n = 10000, https = TRUE)
for(i in 2:length(potentials)){
  potentids <- c(potentids,getPMIDsByKeyWords(journal = potentials[i], dFrom = yr, dTo = yr, n = 10000, https = TRUE))
  Sys.sleep(0.5)
}
lowids <- getPMIDsByKeyWords(journal = lows[1], dFrom = yr, dTo = yr, n = 10000, https = TRUE)
for(i in 2:length(lows)){
  lowids <- c(lowids,getPMIDsByKeyWords(journal = lows[i], dFrom = yr, dTo = yr, n = 10000, https = TRUE))
  Sys.sleep(0.5)
}

highabstracts <- getAbstracts(highids[1])
for (j in 2:length(highids)){
  Sys.sleep(0.25)
  beep_on_error(tryCatch({
    highabstracts <- c(highabstracts,getAbstracts(highids[j]))
  }, error = function(e) {
    Sys.sleep(0.75)
    highabstracts <- c(highabstracts,getAbstracts(highids[j]))
  }), sound = 1)
  #highabstracts <- c(highabstracts,getAbstracts(highids[j]))
  #Sys.sleep(0.2)
  print(j)
  print(Sys.time())
}
beep("complete")

highabstracts <- addpotents(highabstracts)
beep("complete")

lowabstracts <- getAbstracts(lowids[1])
for (j in 2:length(lowids)){
  Sys.sleep(0.25)
  beep_on_error(tryCatch({
    lowabstracts <- c(lowabstracts,getAbstracts(lowids[j]))
  }, error = function(e) {
    Sys.sleep(0.75)
    lowabstracts <- c(lowabstracts,getAbstracts(lowids[j]))
  }), sound = 1)
  #lowabstracts <- c(lowabstracts,getAbstracts(lowids[j]))
  #Sys.sleep(0.2)
  print(j)
  print(Sys.time())
}
beep("fanfare")

abcorp <- Corpus(VectorSource(highabstracts))

abmap <- tm_map(abcorp, PlainTextDocument)
abmap <- tm_map(abmap, removePunctuation)
abmap <- tm_map(abmap, removeWords, c(stopwords('english'),scienceywords))
#abmap <- tm_map(abmap, stemDocument)
# I personally don't like to stem the words

termy <- clean_corpus(abmap)
termy <- breakwords(termy)
term_dtm <- DocumentTermMatrix(termy)
# According to https://www.tidytextmining.com/dtm.html 'Converting to and from...' this is the step
# where we need to convert from corpus into text data using the 'tidy' function

high_words <- tidy(term_dtm)
high_words$tier <- 'High'
high_words$total <- sum(high_words$count)

abcorp <- Corpus(VectorSource(lowabstracts))

abmap <- tm_map(abcorp, PlainTextDocument)
abmap <- tm_map(abmap, removePunctuation)
abmap <- tm_map(abmap, removeWords, stopwords('english'))

termy <- clean_corpus(abmap)
termy <- breakwords(termy)
term_dtm <- DocumentTermMatrix(termy)

low_words <- tidy(term_dtm)
low_words$tier <- 'Low'
low_words$total <- sum(low_words$count)

abstracts <- bind_rows(low_words, high_words)

frequency <- abstracts %>% 
  group_by(tier) %>% 
  count(term, sort = TRUE) %>% 
  left_join(abstracts %>% 
              group_by(tier) %>% 
              summarise(total = n())) %>%
  mutate(freq = n/total)

frequency <- frequency %>% 
  select(tier, term, freq) %>% 
  spread(tier, freq) %>%
  arrange(Low, High)

word_ratios <- abstracts %>%
  count(term, tier) %>%
  group_by(term) %>%
  filter(sum(n) >= 10) %>%
  ungroup() %>%
  spread(tier, n, fill = 0) %>%
  mutate_if(is.numeric, funs((. + 1) / (sum(.) + 1))) %>%
  mutate(logratio = log(High / Low)) %>%
  arrange(desc(logratio))

nam <- paste(as.character(yr),'Contrasts.png')
png(nam, width = 900, height = 450)

word_ratios %>%
  group_by(logratio < 0) %>%
  top_n(15, abs(logratio)) %>%
  ungroup() %>%
  mutate(term = reorder(term, logratio)) %>%
  ggplot(aes(term, logratio, fill = logratio < 0)) +
  geom_col(show.legend = FALSE) +
  coord_flip() +
  ylab("log odds ratio (High/Med)") +
  scale_fill_discrete(name = "", labels = c("High", "Med")) +
  theme_gray(base_size=12*(80/60))

dev.off()

# to view and weed out nonsense words, check dump then add them to sciencywords
dump <- word_ratios %>% group_by(logratio < 0) %>% top_n(50, abs(logratio))

# to look up the odds ratio of a specific term
# word_ratios[match('optogenetics',word_ratios$term),]

nam <- paste(as.character(yr),'Data.RData')
save.image(file = nam)


rm(list = ls())

yr <- 2015
setwd("D:/Box Sync/Research/Meta Review")

highs <- c('Nature Reviews Neuroscience','Nature Neuroscience','Annual Review of Neuroscience','Neuron')
highs <- c(highs,'Molecular Psychiatry','Biological Psychiatry', 'Acta Neuropathologica', 'Progress in Neurobiology', 'Brain')
# 'Trends in Neurosciences' has too short of abstracts

potentials <- c('Nature', 'Science', 'Nature Communications')

keywords = c("brain","brainstem","neuron","neurons","neural","synapse","synaptic","neuroscience","neurotransmitter")
keywords = c(keywords,"amygdala","axonal","dendrite","axons","dendrites","hippocampus","hypothalamus","thalamus")
keywords = c(keywords, "myelination","myelin","cerebellum","sulcus","gyrus")
keywords = c(keywords,"dopamine" , "serotonin" , "gaba" , "glutamate" , "acetylcholine" , "endorphins" , "oxytocin" , "vasopressin")
keywords <- sort(keywords)

lows <- c('Journal of Neurophysiology','Neuroscience','Frontiers in Neuroscience','Brain Structure and Function')
lows <- c(lows,'Brain Research','Neuroscience Bulletin','Brain and Behavior','NeuroReport','European Journal of Neuroscience')
lows <- c(lows,'Brain Research Bulletin','BMC Neuroscience','Neuroscience Research','Neuroscience Letters')

scienceywords = c("may","study","group","effect","effects","increased","decreased","significant")
scienceywords = c(scienceywords, "significantly","higher","lower","compared","control","including","use")
scienceywords = c(scienceywords, "administration","response","groups","different","present","data","one")
scienceywords = c(scienceywords, "two","whether","day","present","reduced","evidence","first","role","right")
scienceywords = c(scienceywords, "elsevier","found","involved","used","suggest","analysis","can","different")
scienceywords = c(scienceywords, "treatment", "associated", "increase", "time", "hours", "differences","activity")
scienceywords = c(scienceywords, "factors", "changes", "respectively", "rights", "trials", "function","mechanisms")
scienceywords = c(scienceywords, "showed", "findings", "min", "days", "well", "received", "however", "also")
scienceywords = c(scienceywords, "observed", "studies", "results", "using", "also", "less", "without", "rate")
scienceywords = c(scienceywords, "high", "level", "research", "measured", "reserved", "patients", "within")
scienceywords = c(scienceywords, "statement", "wiley", "ibro", "sons", "federation", "noteworthy", "periodicals")
scienceywords = c(scienceywords, "john", "societies", "inâ", "isâ", "andâ", "toâ", "abstractcopyright", "etâ", "skill")
scienceywords = c(scienceywords, "extension", "japan", "ireland", "instructed", "presumably", "prepared", "chapter")
scienceywords = c(scienceywords, "reservedthis", "inc", "theâ", "colleagues","incâ", "maximal", "meanwhile", "alternating")
scienceywords = c(scienceywords, "journal", "authors", "ofâ", "comp", "neurol", "american", "imply", "exp", "european")
scienceywords = c(scienceywords, "righthanded", "biology", "press", "commentary", "author", "university", "please")
scienceywords = c(scienceywords, "society", "permissions", "journalspermissionsoupcom", "email", "oxford", "guarantors")
scienceywords = c(scienceywords, "behalf", "examples", "biological", "firestone", "doiaww", "challenges", "inclusions")
scienceywords = c(scienceywords, "morsella", "gane", "experiment","pathwaycopyright","ilî","doiawx","leibovich","institute")
scienceywords = c(scienceywords, "kalisch", "duarte", "lane", "doibrainawy","mahr", "csibra", "psychiatry","advancement")
scienceywords = c(scienceywords, "australian", "ackermann", "shanks", "newell", "bentley", "pepper", "nettle","sciencey")
scienceywords = c(scienceywords, "christiansen", "cop", "copyright", "lake","kline","klines","sci","sweden","over")
scienceywords = c(scienceywords, "entitled","doibrainaww","diseaseâ","science","neurosci")

addpotents <- function(highabstracts)
{
  potentids <- getPMIDsByKeyWords(journal = potentials[1],dFrom = yr,
                                  dTo = yr, n = 10000, https = TRUE )
  for (i in 2:length(potentials)) {
    potentids <- c( potentids, getPMIDsByKeyWords( journal = potentials[i],dFrom = yr,dTo = yr,
                                                   n = 10000, https = TRUE ))
    Sys.sleep(0.5)
  }
  
  for (j in 1:length(potentids)) {
    Sys.sleep(0.25)
    beep_on_error(tryCatch({
      potabstract <- getAbstracts(potentids[j])
    }, error = function(e) {
      Sys.sleep(0.75)
      potabstract <- getAbstracts(potentids[j])
    }), sound = 1)
    
    if (length(potabstract) > 0) {
      if (sum(sapply(keywords, grepl, potabstract)) > 0) {
        highabstracts <- c(highabstracts, potabstract)
      }
    }
    print(j)
    print(Sys.time())
  }
  return(highabstracts)
}

clean_corpus <- function(corpus){
  corpus <- tm_map(corpus, stripWhitespace)
  corpus <- tm_map(corpus, removeNumbers)
  corpus <- tm_map(corpus, removePunctuation)
  corpus <- tm_map(corpus, content_transformer(tolower))
  corpus <- tm_map(corpus, removeWords, c(stopwords("english"),scienceywords))
  return(corpus)
}

breakwords <- function(corpus){
  corpus <- tm_map(corpus, content_transformer(gsub), pattern = "spraguedawley", replacement = "sprague dawley")
  corpus <- tm_map(corpus, content_transformer(gsub), pattern = "attentiondeficithyperactivity", replacement = "attention deficit hyperactivity")
  corpus <- tm_map(corpus, content_transformer(gsub), pattern = "singlenucleotide", replacement = "single nucleotide")
  corpus <- tm_map(corpus, content_transformer(gsub), pattern = "machinelearning", replacement = "machine learning")
  corpus <- tm_map(corpus, content_transformer(gsub), pattern = "genomewide", replacement = "genome wide")
  corpus <- tm_map(corpus, content_transformer(gsub), pattern = "firstepisode", replacement = "first episode")
  corpus <- tm_map(corpus, content_transformer(gsub), pattern = "firstdegree", replacement = "first degree")
  corpus <- tm_map(corpus, content_transformer(gsub), pattern = "obsessivecompulsive", replacement = "obsessive compulsive")
  corpus <- tm_map(corpus, content_transformer(gsub), pattern = "lossoffunction", replacement = "loss of function")
  corpus <- tm_map(corpus, content_transformer(gsub), pattern = "placebocontrolled", replacement = "placebo controlled")
  corpus <- tm_map(corpus, content_transformer(gsub), pattern = "tnfî", replacement = "tnf")
  corpus <- tm_map(corpus, content_transformer(gsub), pattern = "ratscopyright", replacement = "rats copyright")
  corpus <- tm_map(corpus, content_transformer(gsub), pattern = "ischemiareperfusion", replacement = "ischemia repurfusion")
  corpus <- tm_map(corpus, content_transformer(gsub), pattern = "wholeexome", replacement = "whole exome")
  corpus <- tm_map(corpus, content_transformer(gsub), pattern = "patientderived", replacement = "patient derived")
  corpus <- tm_map(corpus, content_transformer(gsub), pattern = "pairedpulse", replacement = "paired pulse")
  corpus <- tm_map(corpus, content_transformer(gsub), pattern = "nearinfrared", replacement = "near infrared")
  corpus <- tm_map(corpus, content_transformer(gsub), pattern = "atrisk", replacement = "at risk")
  corpus <- tm_map(corpus, content_transformer(gsub), pattern = "wholegenome", replacement = "whole genome")
  corpus <- tm_map(corpus, content_transformer(gsub), pattern = "diseaseassociated", replacement = "disease associated")
  corpus <- tm_map(corpus, content_transformer(gsub), pattern = "schizophreniacopyright", replacement = "schizophrenia copyright")
  corpus <- tm_map(corpus, content_transformer(gsub), pattern = "populationbased", replacement = "population based")
  corpus <- tm_map(corpus, content_transformer(gsub), pattern = "celladhesion", replacement = "cell adhesion")
  corpus <- tm_map(corpus, content_transformer(gsub), pattern = "asdcopyright", replacement = "asd copyright")
  corpus <- tm_map(corpus, content_transformer(gsub), pattern = "ligandbinding", replacement = "ligand binding")
  corpus <- tm_map(corpus, content_transformer(gsub), pattern = "overrepresented", replacement = "over represented")
  corpus <- tm_map(corpus, content_transformer(gsub), pattern = "oestrogen", replacement = "estrogen")  
  corpus <- tm_map(corpus, content_transformer(gsub), pattern = "ageing", replacement = "aging")
  corpus <- tm_map(corpus, content_transformer(gsub), pattern = "analog", replacement = "analogue")
  corpus <- tm_map(corpus, content_transformer(gsub), pattern = "tumours", replacement = "tumour")
  corpus <- tm_map(corpus, content_transformer(gsub), pattern = "centres", replacement = "centers")
  corpus <- tm_map(corpus, content_transformer(gsub), pattern = "neighbouring", replacement = "neighboring")
  corpus <- tm_map(corpus, content_transformer(gsub), pattern = "dosedependently", replacement = "dose dependently")
  corpus <- tm_map(corpus, content_transformer(gsub), pattern = "shamoperated", replacement = "sham operated")
  
  #corpus <- tm_map(corpus, content_transformer(gsub), pattern = "spraguedawley", replacement = "sprague")
  return(corpus)
}

highids <- getPMIDsByKeyWords(journal = highs[1], dFrom = yr, dTo = yr, n = 10000, https = TRUE)
for(i in 2:length(highs)){
  highids <- c(highids,getPMIDsByKeyWords(journal = highs[i], dFrom = yr, dTo = yr, n = 10000, https = TRUE))
  Sys.sleep(0.5)
}
potentids <- getPMIDsByKeyWords(journal = potentials[1], dFrom = yr, dTo = yr, n = 10000, https = TRUE)
for(i in 2:length(potentials)){
  potentids <- c(potentids,getPMIDsByKeyWords(journal = potentials[i], dFrom = yr, dTo = yr, n = 10000, https = TRUE))
  Sys.sleep(0.5)
}
lowids <- getPMIDsByKeyWords(journal = lows[1], dFrom = yr, dTo = yr, n = 10000, https = TRUE)
for(i in 2:length(lows)){
  lowids <- c(lowids,getPMIDsByKeyWords(journal = lows[i], dFrom = yr, dTo = yr, n = 10000, https = TRUE))
  Sys.sleep(0.5)
}

highabstracts <- getAbstracts(highids[1])
for (j in 2:length(highids)){
  Sys.sleep(0.25)
  beep_on_error(tryCatch({
    highabstracts <- c(highabstracts,getAbstracts(highids[j]))
  }, error = function(e) {
    Sys.sleep(0.75)
    highabstracts <- c(highabstracts,getAbstracts(highids[j]))
  }), sound = 1)
  #highabstracts <- c(highabstracts,getAbstracts(highids[j]))
  #Sys.sleep(0.2)
  print(j)
  print(Sys.time())
}
beep("complete")

highabstracts <- addpotents(highabstracts)
beep("complete")

lowabstracts <- getAbstracts(lowids[1])
for (j in 2:length(lowids)){
  Sys.sleep(0.25)
  beep_on_error(tryCatch({
    lowabstracts <- c(lowabstracts,getAbstracts(lowids[j]))
  }, error = function(e) {
    Sys.sleep(0.75)
    lowabstracts <- c(lowabstracts,getAbstracts(lowids[j]))
  }), sound = 1)
  #lowabstracts <- c(lowabstracts,getAbstracts(lowids[j]))
  #Sys.sleep(0.2)
  print(j)
  print(Sys.time())
}
beep("fanfare")

abcorp <- Corpus(VectorSource(highabstracts))

abmap <- tm_map(abcorp, PlainTextDocument)
abmap <- tm_map(abmap, removePunctuation)
abmap <- tm_map(abmap, removeWords, c(stopwords('english'),scienceywords))
#abmap <- tm_map(abmap, stemDocument)
# I personally don't like to stem the words

termy <- clean_corpus(abmap)
termy <- breakwords(termy)
term_dtm <- DocumentTermMatrix(termy)
# According to https://www.tidytextmining.com/dtm.html 'Converting to and from...' this is the step
# where we need to convert from corpus into text data using the 'tidy' function

high_words <- tidy(term_dtm)
high_words$tier <- 'High'
high_words$total <- sum(high_words$count)

abcorp <- Corpus(VectorSource(lowabstracts))

abmap <- tm_map(abcorp, PlainTextDocument)
abmap <- tm_map(abmap, removePunctuation)
abmap <- tm_map(abmap, removeWords, stopwords('english'))

termy <- clean_corpus(abmap)
termy <- breakwords(termy)
term_dtm <- DocumentTermMatrix(termy)

low_words <- tidy(term_dtm)
low_words$tier <- 'Low'
low_words$total <- sum(low_words$count)

abstracts <- bind_rows(low_words, high_words)

frequency <- abstracts %>% 
  group_by(tier) %>% 
  count(term, sort = TRUE) %>% 
  left_join(abstracts %>% 
              group_by(tier) %>% 
              summarise(total = n())) %>%
  mutate(freq = n/total)

frequency <- frequency %>% 
  select(tier, term, freq) %>% 
  spread(tier, freq) %>%
  arrange(Low, High)

word_ratios <- abstracts %>%
  count(term, tier) %>%
  group_by(term) %>%
  filter(sum(n) >= 10) %>%
  ungroup() %>%
  spread(tier, n, fill = 0) %>%
  mutate_if(is.numeric, funs((. + 1) / (sum(.) + 1))) %>%
  mutate(logratio = log(High / Low)) %>%
  arrange(desc(logratio))

nam <- paste(as.character(yr),'Contrasts.png')
png(nam, width = 900, height = 450)

word_ratios %>%
  group_by(logratio < 0) %>%
  top_n(15, abs(logratio)) %>%
  ungroup() %>%
  mutate(term = reorder(term, logratio)) %>%
  ggplot(aes(term, logratio, fill = logratio < 0)) +
  geom_col(show.legend = FALSE) +
  coord_flip() +
  ylab("log odds ratio (High/Med)") +
  scale_fill_discrete(name = "", labels = c("High", "Med")) +
  theme_gray(base_size=12*(80/60))

dev.off()

# to view and weed out nonsense words, check dump then add them to sciencywords
dump <- word_ratios %>% group_by(logratio < 0) %>% top_n(50, abs(logratio))

# to look up the odds ratio of a specific term
# word_ratios[match('optogenetics',word_ratios$term),]

nam <- paste(as.character(yr),'Data.RData')
save.image(file = nam)


rm(list = ls())

yr <- 2016
setwd("D:/Box Sync/Research/Meta Review")

highs <- c('Nature Reviews Neuroscience','Nature Neuroscience','Annual Review of Neuroscience','Neuron')
highs <- c(highs,'Molecular Psychiatry','Biological Psychiatry', 'Acta Neuropathologica', 'Progress in Neurobiology', 'Brain')
# 'Trends in Neurosciences' has too short of abstracts

potentials <- c('Nature', 'Science', 'Nature Communications')

keywords = c("brain","brainstem","neuron","neurons","neural","synapse","synaptic","neuroscience","neurotransmitter")
keywords = c(keywords,"amygdala","axonal","dendrite","axons","dendrites","hippocampus","hypothalamus","thalamus")
keywords = c(keywords, "myelination","myelin","cerebellum","sulcus","gyrus")
keywords = c(keywords,"dopamine" , "serotonin" , "gaba" , "glutamate" , "acetylcholine" , "endorphins" , "oxytocin" , "vasopressin")
keywords <- sort(keywords)

lows <- c('Journal of Neurophysiology','Neuroscience','Frontiers in Neuroscience','Brain Structure and Function')
lows <- c(lows,'Brain Research','Neuroscience Bulletin','Brain and Behavior','NeuroReport','European Journal of Neuroscience')
lows <- c(lows,'Brain Research Bulletin','BMC Neuroscience','Neuroscience Research','Neuroscience Letters')

scienceywords = c("may","study","group","effect","effects","increased","decreased","significant")
scienceywords = c(scienceywords, "significantly","higher","lower","compared","control","including","use")
scienceywords = c(scienceywords, "administration","response","groups","different","present","data","one")
scienceywords = c(scienceywords, "two","whether","day","present","reduced","evidence","first","role","right")
scienceywords = c(scienceywords, "elsevier","found","involved","used","suggest","analysis","can","different")
scienceywords = c(scienceywords, "treatment", "associated", "increase", "time", "hours", "differences","activity")
scienceywords = c(scienceywords, "factors", "changes", "respectively", "rights", "trials", "function","mechanisms")
scienceywords = c(scienceywords, "showed", "findings", "min", "days", "well", "received", "however", "also")
scienceywords = c(scienceywords, "observed", "studies", "results", "using", "also", "less", "without", "rate")
scienceywords = c(scienceywords, "high", "level", "research", "measured", "reserved", "patients", "within")
scienceywords = c(scienceywords, "statement", "wiley", "ibro", "sons", "federation", "noteworthy", "periodicals")
scienceywords = c(scienceywords, "john", "societies", "inâ", "isâ", "andâ", "toâ", "abstractcopyright", "etâ", "skill")
scienceywords = c(scienceywords, "extension", "japan", "ireland", "instructed", "presumably", "prepared", "chapter")
scienceywords = c(scienceywords, "reservedthis", "inc", "theâ", "colleagues","incâ", "maximal", "meanwhile", "alternating")
scienceywords = c(scienceywords, "journal", "authors", "ofâ", "comp", "neurol", "american", "imply", "exp", "european")
scienceywords = c(scienceywords, "righthanded", "biology", "press", "commentary", "author", "university", "please")
scienceywords = c(scienceywords, "society", "permissions", "journalspermissionsoupcom", "email", "oxford", "guarantors")
scienceywords = c(scienceywords, "behalf", "examples", "biological", "firestone", "doiaww", "challenges", "inclusions")
scienceywords = c(scienceywords, "morsella", "gane", "experiment","pathwaycopyright","ilî","doiawx","leibovich","institute")
scienceywords = c(scienceywords, "kalisch", "duarte", "lane", "doibrainawy","mahr", "csibra", "psychiatry","advancement")
scienceywords = c(scienceywords, "australian", "ackermann", "shanks", "newell", "bentley", "pepper", "nettle","sciencey")
scienceywords = c(scienceywords, "christiansen", "cop", "copyright", "lake","kline","klines","sci","sweden","over")
scienceywords = c(scienceywords, "entitled","doibrainaww","diseaseâ","science","neurosci")

addpotents <- function(highabstracts)
{
  potentids <- getPMIDsByKeyWords(journal = potentials[1],dFrom = yr,
                                  dTo = yr, n = 10000, https = TRUE )
  for (i in 2:length(potentials)) {
    potentids <- c( potentids, getPMIDsByKeyWords( journal = potentials[i],dFrom = yr,dTo = yr,
                                                   n = 10000, https = TRUE ))
    Sys.sleep(0.5)
  }
  
  for (j in 1:length(potentids)) {
    Sys.sleep(0.25)
    beep_on_error(tryCatch({
      potabstract <- getAbstracts(potentids[j])
    }, error = function(e) {
      Sys.sleep(0.75)
      potabstract <- getAbstracts(potentids[j])
    }), sound = 1)
    
    if (length(potabstract) > 0) {
      if (sum(sapply(keywords, grepl, potabstract)) > 0) {
        highabstracts <- c(highabstracts, potabstract)
      }
    }
    print(j)
    print(Sys.time())
  }
  return(highabstracts)
}

clean_corpus <- function(corpus){
  corpus <- tm_map(corpus, stripWhitespace)
  corpus <- tm_map(corpus, removeNumbers)
  corpus <- tm_map(corpus, removePunctuation)
  corpus <- tm_map(corpus, content_transformer(tolower))
  corpus <- tm_map(corpus, removeWords, c(stopwords("english"),scienceywords))
  return(corpus)
}

breakwords <- function(corpus){
  corpus <- tm_map(corpus, content_transformer(gsub), pattern = "spraguedawley", replacement = "sprague dawley")
  corpus <- tm_map(corpus, content_transformer(gsub), pattern = "attentiondeficithyperactivity", replacement = "attention deficit hyperactivity")
  corpus <- tm_map(corpus, content_transformer(gsub), pattern = "singlenucleotide", replacement = "single nucleotide")
  corpus <- tm_map(corpus, content_transformer(gsub), pattern = "machinelearning", replacement = "machine learning")
  corpus <- tm_map(corpus, content_transformer(gsub), pattern = "genomewide", replacement = "genome wide")
  corpus <- tm_map(corpus, content_transformer(gsub), pattern = "firstepisode", replacement = "first episode")
  corpus <- tm_map(corpus, content_transformer(gsub), pattern = "firstdegree", replacement = "first degree")
  corpus <- tm_map(corpus, content_transformer(gsub), pattern = "obsessivecompulsive", replacement = "obsessive compulsive")
  corpus <- tm_map(corpus, content_transformer(gsub), pattern = "lossoffunction", replacement = "loss of function")
  corpus <- tm_map(corpus, content_transformer(gsub), pattern = "placebocontrolled", replacement = "placebo controlled")
  corpus <- tm_map(corpus, content_transformer(gsub), pattern = "tnfî", replacement = "tnf")
  corpus <- tm_map(corpus, content_transformer(gsub), pattern = "ratscopyright", replacement = "rats copyright")
  corpus <- tm_map(corpus, content_transformer(gsub), pattern = "ischemiareperfusion", replacement = "ischemia repurfusion")
  corpus <- tm_map(corpus, content_transformer(gsub), pattern = "wholeexome", replacement = "whole exome")
  corpus <- tm_map(corpus, content_transformer(gsub), pattern = "patientderived", replacement = "patient derived")
  corpus <- tm_map(corpus, content_transformer(gsub), pattern = "pairedpulse", replacement = "paired pulse")
  corpus <- tm_map(corpus, content_transformer(gsub), pattern = "nearinfrared", replacement = "near infrared")
  corpus <- tm_map(corpus, content_transformer(gsub), pattern = "atrisk", replacement = "at risk")
  corpus <- tm_map(corpus, content_transformer(gsub), pattern = "wholegenome", replacement = "whole genome")
  corpus <- tm_map(corpus, content_transformer(gsub), pattern = "diseaseassociated", replacement = "disease associated")
  corpus <- tm_map(corpus, content_transformer(gsub), pattern = "schizophreniacopyright", replacement = "schizophrenia copyright")
  corpus <- tm_map(corpus, content_transformer(gsub), pattern = "populationbased", replacement = "population based")
  corpus <- tm_map(corpus, content_transformer(gsub), pattern = "celladhesion", replacement = "cell adhesion")
  corpus <- tm_map(corpus, content_transformer(gsub), pattern = "asdcopyright", replacement = "asd copyright")
  corpus <- tm_map(corpus, content_transformer(gsub), pattern = "ligandbinding", replacement = "ligand binding")
  corpus <- tm_map(corpus, content_transformer(gsub), pattern = "overrepresented", replacement = "over represented")
  corpus <- tm_map(corpus, content_transformer(gsub), pattern = "oestrogen", replacement = "estrogen")  
  corpus <- tm_map(corpus, content_transformer(gsub), pattern = "ageing", replacement = "aging")
  corpus <- tm_map(corpus, content_transformer(gsub), pattern = "analog", replacement = "analogue")
  corpus <- tm_map(corpus, content_transformer(gsub), pattern = "tumours", replacement = "tumour")
  corpus <- tm_map(corpus, content_transformer(gsub), pattern = "centres", replacement = "centers")
  corpus <- tm_map(corpus, content_transformer(gsub), pattern = "neighbouring", replacement = "neighboring")
  corpus <- tm_map(corpus, content_transformer(gsub), pattern = "dosedependently", replacement = "dose dependently")
  corpus <- tm_map(corpus, content_transformer(gsub), pattern = "shamoperated", replacement = "sham operated")
  
  #corpus <- tm_map(corpus, content_transformer(gsub), pattern = "spraguedawley", replacement = "sprague")
  return(corpus)
}

highids <- getPMIDsByKeyWords(journal = highs[1], dFrom = yr, dTo = yr, n = 10000, https = TRUE)
for(i in 2:length(highs)){
  highids <- c(highids,getPMIDsByKeyWords(journal = highs[i], dFrom = yr, dTo = yr, n = 10000, https = TRUE))
  Sys.sleep(0.5)
}
potentids <- getPMIDsByKeyWords(journal = potentials[1], dFrom = yr, dTo = yr, n = 10000, https = TRUE)
for(i in 2:length(potentials)){
  potentids <- c(potentids,getPMIDsByKeyWords(journal = potentials[i], dFrom = yr, dTo = yr, n = 10000, https = TRUE))
  Sys.sleep(0.5)
}
lowids <- getPMIDsByKeyWords(journal = lows[1], dFrom = yr, dTo = yr, n = 10000, https = TRUE)
for(i in 2:length(lows)){
  lowids <- c(lowids,getPMIDsByKeyWords(journal = lows[i], dFrom = yr, dTo = yr, n = 10000, https = TRUE))
  Sys.sleep(0.5)
}

highabstracts <- getAbstracts(highids[1])
for (j in 2:length(highids)){
  Sys.sleep(0.25)
  beep_on_error(tryCatch({
    highabstracts <- c(highabstracts,getAbstracts(highids[j]))
  }, error = function(e) {
    Sys.sleep(0.75)
    highabstracts <- c(highabstracts,getAbstracts(highids[j]))
  }), sound = 1)
  #highabstracts <- c(highabstracts,getAbstracts(highids[j]))
  #Sys.sleep(0.2)
  print(j)
  print(Sys.time())
}
beep("complete")

highabstracts <- addpotents(highabstracts)
beep("complete")

lowabstracts <- getAbstracts(lowids[1])
for (j in 2:length(lowids)){
  Sys.sleep(0.25)
  beep_on_error(tryCatch({
    lowabstracts <- c(lowabstracts,getAbstracts(lowids[j]))
  }, error = function(e) {
    Sys.sleep(0.75)
    lowabstracts <- c(lowabstracts,getAbstracts(lowids[j]))
  }), sound = 1)
  #lowabstracts <- c(lowabstracts,getAbstracts(lowids[j]))
  #Sys.sleep(0.2)
  print(j)
  print(Sys.time())
}
beep("fanfare")

abcorp <- Corpus(VectorSource(highabstracts))

abmap <- tm_map(abcorp, PlainTextDocument)
abmap <- tm_map(abmap, removePunctuation)
abmap <- tm_map(abmap, removeWords, c(stopwords('english'),scienceywords))
#abmap <- tm_map(abmap, stemDocument)
# I personally don't like to stem the words

termy <- clean_corpus(abmap)
termy <- breakwords(termy)
term_dtm <- DocumentTermMatrix(termy)
# According to https://www.tidytextmining.com/dtm.html 'Converting to and from...' this is the step
# where we need to convert from corpus into text data using the 'tidy' function

high_words <- tidy(term_dtm)
high_words$tier <- 'High'
high_words$total <- sum(high_words$count)

abcorp <- Corpus(VectorSource(lowabstracts))

abmap <- tm_map(abcorp, PlainTextDocument)
abmap <- tm_map(abmap, removePunctuation)
abmap <- tm_map(abmap, removeWords, stopwords('english'))

termy <- clean_corpus(abmap)
termy <- breakwords(termy)
term_dtm <- DocumentTermMatrix(termy)

low_words <- tidy(term_dtm)
low_words$tier <- 'Low'
low_words$total <- sum(low_words$count)

abstracts <- bind_rows(low_words, high_words)

frequency <- abstracts %>% 
  group_by(tier) %>% 
  count(term, sort = TRUE) %>% 
  left_join(abstracts %>% 
              group_by(tier) %>% 
              summarise(total = n())) %>%
  mutate(freq = n/total)

frequency <- frequency %>% 
  select(tier, term, freq) %>% 
  spread(tier, freq) %>%
  arrange(Low, High)

word_ratios <- abstracts %>%
  count(term, tier) %>%
  group_by(term) %>%
  filter(sum(n) >= 10) %>%
  ungroup() %>%
  spread(tier, n, fill = 0) %>%
  mutate_if(is.numeric, funs((. + 1) / (sum(.) + 1))) %>%
  mutate(logratio = log(High / Low)) %>%
  arrange(desc(logratio))

nam <- paste(as.character(yr),'Contrasts.png')
png(nam, width = 900, height = 450)

word_ratios %>%
  group_by(logratio < 0) %>%
  top_n(15, abs(logratio)) %>%
  ungroup() %>%
  mutate(term = reorder(term, logratio)) %>%
  ggplot(aes(term, logratio, fill = logratio < 0)) +
  geom_col(show.legend = FALSE) +
  coord_flip() +
  ylab("log odds ratio (High/Med)") +
  scale_fill_discrete(name = "", labels = c("High", "Med")) +
  theme_gray(base_size=12*(80/60))

dev.off()

# to view and weed out nonsense words, check dump then add them to sciencywords
dump <- word_ratios %>% group_by(logratio < 0) %>% top_n(50, abs(logratio))

# to look up the odds ratio of a specific term
# word_ratios[match('optogenetics',word_ratios$term),]

nam <- paste(as.character(yr),'Data.RData')
save.image(file = nam)


rm(list = ls())

yr <- 2017
setwd("D:/Box Sync/Research/Meta Review")

highs <- c('Nature Reviews Neuroscience','Nature Neuroscience','Annual Review of Neuroscience','Neuron')
highs <- c(highs,'Molecular Psychiatry','Biological Psychiatry', 'Acta Neuropathologica', 'Progress in Neurobiology', 'Brain')
# 'Trends in Neurosciences' has too short of abstracts

potentials <- c('Nature', 'Science', 'Nature Communications')

keywords = c("brain","brainstem","neuron","neurons","neural","synapse","synaptic","neuroscience","neurotransmitter")
keywords = c(keywords,"amygdala","axonal","dendrite","axons","dendrites","hippocampus","hypothalamus","thalamus")
keywords = c(keywords, "myelination","myelin","cerebellum","sulcus","gyrus")
keywords = c(keywords,"dopamine" , "serotonin" , "gaba" , "glutamate" , "acetylcholine" , "endorphins" , "oxytocin" , "vasopressin")
keywords <- sort(keywords)

lows <- c('Journal of Neurophysiology','Neuroscience','Frontiers in Neuroscience','Brain Structure and Function')
lows <- c(lows,'Brain Research','Neuroscience Bulletin','Brain and Behavior','NeuroReport','European Journal of Neuroscience')
lows <- c(lows,'Brain Research Bulletin','BMC Neuroscience','Neuroscience Research','Neuroscience Letters')

scienceywords = c("may","study","group","effect","effects","increased","decreased","significant")
scienceywords = c(scienceywords, "significantly","higher","lower","compared","control","including","use")
scienceywords = c(scienceywords, "administration","response","groups","different","present","data","one")
scienceywords = c(scienceywords, "two","whether","day","present","reduced","evidence","first","role","right")
scienceywords = c(scienceywords, "elsevier","found","involved","used","suggest","analysis","can","different")
scienceywords = c(scienceywords, "treatment", "associated", "increase", "time", "hours", "differences","activity")
scienceywords = c(scienceywords, "factors", "changes", "respectively", "rights", "trials", "function","mechanisms")
scienceywords = c(scienceywords, "showed", "findings", "min", "days", "well", "received", "however", "also")
scienceywords = c(scienceywords, "observed", "studies", "results", "using", "also", "less", "without", "rate")
scienceywords = c(scienceywords, "high", "level", "research", "measured", "reserved", "patients", "within")
scienceywords = c(scienceywords, "statement", "wiley", "ibro", "sons", "federation", "noteworthy", "periodicals")
scienceywords = c(scienceywords, "john", "societies", "inâ", "isâ", "andâ", "toâ", "abstractcopyright", "etâ", "skill")
scienceywords = c(scienceywords, "extension", "japan", "ireland", "instructed", "presumably", "prepared", "chapter")
scienceywords = c(scienceywords, "reservedthis", "inc", "theâ", "colleagues","incâ", "maximal", "meanwhile", "alternating")
scienceywords = c(scienceywords, "journal", "authors", "ofâ", "comp", "neurol", "american", "imply", "exp", "european")
scienceywords = c(scienceywords, "righthanded", "biology", "press", "commentary", "author", "university", "please")
scienceywords = c(scienceywords, "society", "permissions", "journalspermissionsoupcom", "email", "oxford", "guarantors")
scienceywords = c(scienceywords, "behalf", "examples", "biological", "firestone", "doiaww", "challenges", "inclusions")
scienceywords = c(scienceywords, "morsella", "gane", "experiment","pathwaycopyright","ilî","doiawx","leibovich","institute")
scienceywords = c(scienceywords, "kalisch", "duarte", "lane", "doibrainawy","mahr", "csibra", "psychiatry","advancement")
scienceywords = c(scienceywords, "australian", "ackermann", "shanks", "newell", "bentley", "pepper", "nettle","sciencey")
scienceywords = c(scienceywords, "christiansen", "cop", "copyright", "lake","kline","klines","sci","sweden","over")
scienceywords = c(scienceywords, "entitled","doibrainaww","diseaseâ","science","neurosci")

addpotents <- function(highabstracts)
{
  potentids <- getPMIDsByKeyWords(journal = potentials[1],dFrom = yr,
                                  dTo = yr, n = 10000, https = TRUE )
  for (i in 2:length(potentials)) {
    potentids <- c( potentids, getPMIDsByKeyWords( journal = potentials[i],dFrom = yr,dTo = yr,
                                                   n = 10000, https = TRUE ))
    Sys.sleep(0.5)
  }
  
  for (j in 1:length(potentids)) {
    Sys.sleep(0.25)
    beep_on_error(tryCatch({
      potabstract <- getAbstracts(potentids[j])
    }, error = function(e) {
      Sys.sleep(0.75)
      potabstract <- getAbstracts(potentids[j])
    }), sound = 1)
    
    if (length(potabstract) > 0) {
      if (sum(sapply(keywords, grepl, potabstract)) > 0) {
        highabstracts <- c(highabstracts, potabstract)
      }
    }
    print(j)
    print(Sys.time())
  }
  return(highabstracts)
}

clean_corpus <- function(corpus){
  corpus <- tm_map(corpus, stripWhitespace)
  corpus <- tm_map(corpus, removeNumbers)
  corpus <- tm_map(corpus, removePunctuation)
  corpus <- tm_map(corpus, content_transformer(tolower))
  corpus <- tm_map(corpus, removeWords, c(stopwords("english"),scienceywords))
  return(corpus)
}

breakwords <- function(corpus){
  corpus <- tm_map(corpus, content_transformer(gsub), pattern = "spraguedawley", replacement = "sprague dawley")
  corpus <- tm_map(corpus, content_transformer(gsub), pattern = "attentiondeficithyperactivity", replacement = "attention deficit hyperactivity")
  corpus <- tm_map(corpus, content_transformer(gsub), pattern = "singlenucleotide", replacement = "single nucleotide")
  corpus <- tm_map(corpus, content_transformer(gsub), pattern = "machinelearning", replacement = "machine learning")
  corpus <- tm_map(corpus, content_transformer(gsub), pattern = "genomewide", replacement = "genome wide")
  corpus <- tm_map(corpus, content_transformer(gsub), pattern = "firstepisode", replacement = "first episode")
  corpus <- tm_map(corpus, content_transformer(gsub), pattern = "firstdegree", replacement = "first degree")
  corpus <- tm_map(corpus, content_transformer(gsub), pattern = "obsessivecompulsive", replacement = "obsessive compulsive")
  corpus <- tm_map(corpus, content_transformer(gsub), pattern = "lossoffunction", replacement = "loss of function")
  corpus <- tm_map(corpus, content_transformer(gsub), pattern = "placebocontrolled", replacement = "placebo controlled")
  corpus <- tm_map(corpus, content_transformer(gsub), pattern = "tnfî", replacement = "tnf")
  corpus <- tm_map(corpus, content_transformer(gsub), pattern = "ratscopyright", replacement = "rats copyright")
  corpus <- tm_map(corpus, content_transformer(gsub), pattern = "ischemiareperfusion", replacement = "ischemia repurfusion")
  corpus <- tm_map(corpus, content_transformer(gsub), pattern = "wholeexome", replacement = "whole exome")
  corpus <- tm_map(corpus, content_transformer(gsub), pattern = "patientderived", replacement = "patient derived")
  corpus <- tm_map(corpus, content_transformer(gsub), pattern = "pairedpulse", replacement = "paired pulse")
  corpus <- tm_map(corpus, content_transformer(gsub), pattern = "nearinfrared", replacement = "near infrared")
  corpus <- tm_map(corpus, content_transformer(gsub), pattern = "atrisk", replacement = "at risk")
  corpus <- tm_map(corpus, content_transformer(gsub), pattern = "wholegenome", replacement = "whole genome")
  corpus <- tm_map(corpus, content_transformer(gsub), pattern = "diseaseassociated", replacement = "disease associated")
  corpus <- tm_map(corpus, content_transformer(gsub), pattern = "schizophreniacopyright", replacement = "schizophrenia copyright")
  corpus <- tm_map(corpus, content_transformer(gsub), pattern = "populationbased", replacement = "population based")
  corpus <- tm_map(corpus, content_transformer(gsub), pattern = "celladhesion", replacement = "cell adhesion")
  corpus <- tm_map(corpus, content_transformer(gsub), pattern = "asdcopyright", replacement = "asd copyright")
  corpus <- tm_map(corpus, content_transformer(gsub), pattern = "ligandbinding", replacement = "ligand binding")
  corpus <- tm_map(corpus, content_transformer(gsub), pattern = "overrepresented", replacement = "over represented")
  corpus <- tm_map(corpus, content_transformer(gsub), pattern = "oestrogen", replacement = "estrogen")  
  corpus <- tm_map(corpus, content_transformer(gsub), pattern = "ageing", replacement = "aging")
  corpus <- tm_map(corpus, content_transformer(gsub), pattern = "analog", replacement = "analogue")
  corpus <- tm_map(corpus, content_transformer(gsub), pattern = "tumours", replacement = "tumour")
  corpus <- tm_map(corpus, content_transformer(gsub), pattern = "centres", replacement = "centers")
  corpus <- tm_map(corpus, content_transformer(gsub), pattern = "neighbouring", replacement = "neighboring")
  corpus <- tm_map(corpus, content_transformer(gsub), pattern = "dosedependently", replacement = "dose dependently")
  corpus <- tm_map(corpus, content_transformer(gsub), pattern = "shamoperated", replacement = "sham operated")
  
  #corpus <- tm_map(corpus, content_transformer(gsub), pattern = "spraguedawley", replacement = "sprague")
  return(corpus)
}

highids <- getPMIDsByKeyWords(journal = highs[1], dFrom = yr, dTo = yr, n = 10000, https = TRUE)
for(i in 2:length(highs)){
  highids <- c(highids,getPMIDsByKeyWords(journal = highs[i], dFrom = yr, dTo = yr, n = 10000, https = TRUE))
  Sys.sleep(0.5)
}
potentids <- getPMIDsByKeyWords(journal = potentials[1], dFrom = yr, dTo = yr, n = 10000, https = TRUE)
for(i in 2:length(potentials)){
  potentids <- c(potentids,getPMIDsByKeyWords(journal = potentials[i], dFrom = yr, dTo = yr, n = 10000, https = TRUE))
  Sys.sleep(0.5)
}
lowids <- getPMIDsByKeyWords(journal = lows[1], dFrom = yr, dTo = yr, n = 10000, https = TRUE)
for(i in 2:length(lows)){
  lowids <- c(lowids,getPMIDsByKeyWords(journal = lows[i], dFrom = yr, dTo = yr, n = 10000, https = TRUE))
  Sys.sleep(0.5)
}

highabstracts <- getAbstracts(highids[1])
for (j in 2:length(highids)){
  Sys.sleep(0.25)
  beep_on_error(tryCatch({
    highabstracts <- c(highabstracts,getAbstracts(highids[j]))
  }, error = function(e) {
    Sys.sleep(0.75)
    highabstracts <- c(highabstracts,getAbstracts(highids[j]))
  }), sound = 1)
  #highabstracts <- c(highabstracts,getAbstracts(highids[j]))
  #Sys.sleep(0.2)
  print(j)
  print(Sys.time())
}
beep("complete")

highabstracts <- addpotents(highabstracts)
beep("complete")

lowabstracts <- getAbstracts(lowids[1])
for (j in 2:length(lowids)){
  Sys.sleep(0.25)
  beep_on_error(tryCatch({
    lowabstracts <- c(lowabstracts,getAbstracts(lowids[j]))
  }, error = function(e) {
    Sys.sleep(0.75)
    lowabstracts <- c(lowabstracts,getAbstracts(lowids[j]))
  }), sound = 1)
  #lowabstracts <- c(lowabstracts,getAbstracts(lowids[j]))
  #Sys.sleep(0.2)
  print(j)
  print(Sys.time())
}
beep("fanfare")

abcorp <- Corpus(VectorSource(highabstracts))

abmap <- tm_map(abcorp, PlainTextDocument)
abmap <- tm_map(abmap, removePunctuation)
abmap <- tm_map(abmap, removeWords, c(stopwords('english'),scienceywords))
#abmap <- tm_map(abmap, stemDocument)
# I personally don't like to stem the words

termy <- clean_corpus(abmap)
termy <- breakwords(termy)
term_dtm <- DocumentTermMatrix(termy)
# According to https://www.tidytextmining.com/dtm.html 'Converting to and from...' this is the step
# where we need to convert from corpus into text data using the 'tidy' function

high_words <- tidy(term_dtm)
high_words$tier <- 'High'
high_words$total <- sum(high_words$count)

abcorp <- Corpus(VectorSource(lowabstracts))

abmap <- tm_map(abcorp, PlainTextDocument)
abmap <- tm_map(abmap, removePunctuation)
abmap <- tm_map(abmap, removeWords, stopwords('english'))

termy <- clean_corpus(abmap)
termy <- breakwords(termy)
term_dtm <- DocumentTermMatrix(termy)

low_words <- tidy(term_dtm)
low_words$tier <- 'Low'
low_words$total <- sum(low_words$count)

abstracts <- bind_rows(low_words, high_words)

frequency <- abstracts %>% 
  group_by(tier) %>% 
  count(term, sort = TRUE) %>% 
  left_join(abstracts %>% 
              group_by(tier) %>% 
              summarise(total = n())) %>%
  mutate(freq = n/total)

frequency <- frequency %>% 
  select(tier, term, freq) %>% 
  spread(tier, freq) %>%
  arrange(Low, High)

word_ratios <- abstracts %>%
  count(term, tier) %>%
  group_by(term) %>%
  filter(sum(n) >= 10) %>%
  ungroup() %>%
  spread(tier, n, fill = 0) %>%
  mutate_if(is.numeric, funs((. + 1) / (sum(.) + 1))) %>%
  mutate(logratio = log(High / Low)) %>%
  arrange(desc(logratio))

nam <- paste(as.character(yr),'Contrasts.png')
png(nam, width = 900, height = 450)

word_ratios %>%
  group_by(logratio < 0) %>%
  top_n(15, abs(logratio)) %>%
  ungroup() %>%
  mutate(term = reorder(term, logratio)) %>%
  ggplot(aes(term, logratio, fill = logratio < 0)) +
  geom_col(show.legend = FALSE) +
  coord_flip() +
  ylab("log odds ratio (High/Med)") +
  scale_fill_discrete(name = "", labels = c("High", "Med")) +
  theme_gray(base_size=12*(80/60))

dev.off()

# to view and weed out nonsense words, check dump then add them to sciencywords
dump <- word_ratios %>% group_by(logratio < 0) %>% top_n(50, abs(logratio))

# to look up the odds ratio of a specific term
# word_ratios[match('optogenetics',word_ratios$term),]

nam <- paste(as.character(yr),'Data.RData')
save.image(file = nam)


rm(list = ls())

yr <- 2018
setwd("D:/Box Sync/Research/Meta Review")

highs <- c('Nature Reviews Neuroscience','Nature Neuroscience','Annual Review of Neuroscience','Neuron')
highs <- c(highs,'Molecular Psychiatry','Biological Psychiatry', 'Acta Neuropathologica', 'Progress in Neurobiology', 'Brain')
# 'Trends in Neurosciences' has too short of abstracts

potentials <- c('Nature', 'Science', 'Nature Communications')

keywords = c("brain","brainstem","neuron","neurons","neural","synapse","synaptic","neuroscience","neurotransmitter")
keywords = c(keywords,"amygdala","axonal","dendrite","axons","dendrites","hippocampus","hypothalamus","thalamus")
keywords = c(keywords, "myelination","myelin","cerebellum","sulcus","gyrus")
keywords = c(keywords,"dopamine" , "serotonin" , "gaba" , "glutamate" , "acetylcholine" , "endorphins" , "oxytocin" , "vasopressin")
keywords <- sort(keywords)

lows <- c('Journal of Neurophysiology','Neuroscience','Frontiers in Neuroscience','Brain Structure and Function')
lows <- c(lows,'Brain Research','Neuroscience Bulletin','Brain and Behavior','NeuroReport','European Journal of Neuroscience')
lows <- c(lows,'Brain Research Bulletin','BMC Neuroscience','Neuroscience Research','Neuroscience Letters')

scienceywords = c("may","study","group","effect","effects","increased","decreased","significant")
scienceywords = c(scienceywords, "significantly","higher","lower","compared","control","including","use")
scienceywords = c(scienceywords, "administration","response","groups","different","present","data","one")
scienceywords = c(scienceywords, "two","whether","day","present","reduced","evidence","first","role","right")
scienceywords = c(scienceywords, "elsevier","found","involved","used","suggest","analysis","can","different")
scienceywords = c(scienceywords, "treatment", "associated", "increase", "time", "hours", "differences","activity")
scienceywords = c(scienceywords, "factors", "changes", "respectively", "rights", "trials", "function","mechanisms")
scienceywords = c(scienceywords, "showed", "findings", "min", "days", "well", "received", "however", "also")
scienceywords = c(scienceywords, "observed", "studies", "results", "using", "also", "less", "without", "rate")
scienceywords = c(scienceywords, "high", "level", "research", "measured", "reserved", "patients", "within")
scienceywords = c(scienceywords, "statement", "wiley", "ibro", "sons", "federation", "noteworthy", "periodicals")
scienceywords = c(scienceywords, "john", "societies", "inâ", "isâ", "andâ", "toâ", "abstractcopyright", "etâ", "skill")
scienceywords = c(scienceywords, "extension", "japan", "ireland", "instructed", "presumably", "prepared", "chapter")
scienceywords = c(scienceywords, "reservedthis", "inc", "theâ", "colleagues","incâ", "maximal", "meanwhile", "alternating")
scienceywords = c(scienceywords, "journal", "authors", "ofâ", "comp", "neurol", "american", "imply", "exp", "european")
scienceywords = c(scienceywords, "righthanded", "biology", "press", "commentary", "author", "university", "please")
scienceywords = c(scienceywords, "society", "permissions", "journalspermissionsoupcom", "email", "oxford", "guarantors")
scienceywords = c(scienceywords, "behalf", "examples", "biological", "firestone", "doiaww", "challenges", "inclusions")
scienceywords = c(scienceywords, "morsella", "gane", "experiment","pathwaycopyright","ilî","doiawx","leibovich","institute")
scienceywords = c(scienceywords, "kalisch", "duarte", "lane", "doibrainawy","mahr", "csibra", "psychiatry","advancement")
scienceywords = c(scienceywords, "australian", "ackermann", "shanks", "newell", "bentley", "pepper", "nettle","sciencey")
scienceywords = c(scienceywords, "christiansen", "cop", "copyright", "lake","kline","klines","sci","sweden","over")
scienceywords = c(scienceywords, "entitled","doibrainaww","diseaseâ","science","neurosci")

addpotents <- function(highabstracts)
{
  potentids <- getPMIDsByKeyWords(journal = potentials[1],dFrom = yr,
                                  dTo = yr, n = 10000, https = TRUE )
  for (i in 2:length(potentials)) {
    potentids <- c( potentids, getPMIDsByKeyWords( journal = potentials[i],dFrom = yr,dTo = yr,
                                                   n = 10000, https = TRUE ))
    Sys.sleep(0.5)
  }
  
  for (j in 1:length(potentids)) {
    Sys.sleep(0.25)
    beep_on_error(tryCatch({
      potabstract <- getAbstracts(potentids[j])
    }, error = function(e) {
      Sys.sleep(0.75)
      potabstract <- getAbstracts(potentids[j])
    }), sound = 1)
    
    if (length(potabstract) > 0) {
      if (sum(sapply(keywords, grepl, potabstract)) > 0) {
        highabstracts <- c(highabstracts, potabstract)
      }
    }
    print(j)
    print(Sys.time())
  }
  return(highabstracts)
}

clean_corpus <- function(corpus){
  corpus <- tm_map(corpus, stripWhitespace)
  corpus <- tm_map(corpus, removeNumbers)
  corpus <- tm_map(corpus, removePunctuation)
  corpus <- tm_map(corpus, content_transformer(tolower))
  corpus <- tm_map(corpus, removeWords, c(stopwords("english"),scienceywords))
  return(corpus)
}

breakwords <- function(corpus){
  corpus <- tm_map(corpus, content_transformer(gsub), pattern = "spraguedawley", replacement = "sprague dawley")
  corpus <- tm_map(corpus, content_transformer(gsub), pattern = "attentiondeficithyperactivity", replacement = "attention deficit hyperactivity")
  corpus <- tm_map(corpus, content_transformer(gsub), pattern = "singlenucleotide", replacement = "single nucleotide")
  corpus <- tm_map(corpus, content_transformer(gsub), pattern = "machinelearning", replacement = "machine learning")
  corpus <- tm_map(corpus, content_transformer(gsub), pattern = "genomewide", replacement = "genome wide")
  corpus <- tm_map(corpus, content_transformer(gsub), pattern = "firstepisode", replacement = "first episode")
  corpus <- tm_map(corpus, content_transformer(gsub), pattern = "firstdegree", replacement = "first degree")
  corpus <- tm_map(corpus, content_transformer(gsub), pattern = "obsessivecompulsive", replacement = "obsessive compulsive")
  corpus <- tm_map(corpus, content_transformer(gsub), pattern = "lossoffunction", replacement = "loss of function")
  corpus <- tm_map(corpus, content_transformer(gsub), pattern = "placebocontrolled", replacement = "placebo controlled")
  corpus <- tm_map(corpus, content_transformer(gsub), pattern = "tnfî", replacement = "tnf")
  corpus <- tm_map(corpus, content_transformer(gsub), pattern = "ratscopyright", replacement = "rats copyright")
  corpus <- tm_map(corpus, content_transformer(gsub), pattern = "ischemiareperfusion", replacement = "ischemia repurfusion")
  corpus <- tm_map(corpus, content_transformer(gsub), pattern = "wholeexome", replacement = "whole exome")
  corpus <- tm_map(corpus, content_transformer(gsub), pattern = "patientderived", replacement = "patient derived")
  corpus <- tm_map(corpus, content_transformer(gsub), pattern = "pairedpulse", replacement = "paired pulse")
  corpus <- tm_map(corpus, content_transformer(gsub), pattern = "nearinfrared", replacement = "near infrared")
  corpus <- tm_map(corpus, content_transformer(gsub), pattern = "atrisk", replacement = "at risk")
  corpus <- tm_map(corpus, content_transformer(gsub), pattern = "wholegenome", replacement = "whole genome")
  corpus <- tm_map(corpus, content_transformer(gsub), pattern = "diseaseassociated", replacement = "disease associated")
  corpus <- tm_map(corpus, content_transformer(gsub), pattern = "schizophreniacopyright", replacement = "schizophrenia copyright")
  corpus <- tm_map(corpus, content_transformer(gsub), pattern = "populationbased", replacement = "population based")
  corpus <- tm_map(corpus, content_transformer(gsub), pattern = "celladhesion", replacement = "cell adhesion")
  corpus <- tm_map(corpus, content_transformer(gsub), pattern = "asdcopyright", replacement = "asd copyright")
  corpus <- tm_map(corpus, content_transformer(gsub), pattern = "ligandbinding", replacement = "ligand binding")
  corpus <- tm_map(corpus, content_transformer(gsub), pattern = "overrepresented", replacement = "over represented")
  corpus <- tm_map(corpus, content_transformer(gsub), pattern = "oestrogen", replacement = "estrogen")  
  corpus <- tm_map(corpus, content_transformer(gsub), pattern = "ageing", replacement = "aging")
  corpus <- tm_map(corpus, content_transformer(gsub), pattern = "analog", replacement = "analogue")
  corpus <- tm_map(corpus, content_transformer(gsub), pattern = "tumours", replacement = "tumour")
  corpus <- tm_map(corpus, content_transformer(gsub), pattern = "centres", replacement = "centers")
  corpus <- tm_map(corpus, content_transformer(gsub), pattern = "neighbouring", replacement = "neighboring")
  corpus <- tm_map(corpus, content_transformer(gsub), pattern = "dosedependently", replacement = "dose dependently")
  corpus <- tm_map(corpus, content_transformer(gsub), pattern = "shamoperated", replacement = "sham operated")
  
  #corpus <- tm_map(corpus, content_transformer(gsub), pattern = "spraguedawley", replacement = "sprague")
  return(corpus)
}

highids <- getPMIDsByKeyWords(journal = highs[1], dFrom = yr, dTo = yr, n = 10000, https = TRUE)
for(i in 2:length(highs)){
  highids <- c(highids,getPMIDsByKeyWords(journal = highs[i], dFrom = yr, dTo = yr, n = 10000, https = TRUE))
  Sys.sleep(0.5)
}
potentids <- getPMIDsByKeyWords(journal = potentials[1], dFrom = yr, dTo = yr, n = 10000, https = TRUE)
for(i in 2:length(potentials)){
  potentids <- c(potentids,getPMIDsByKeyWords(journal = potentials[i], dFrom = yr, dTo = yr, n = 10000, https = TRUE))
  Sys.sleep(0.5)
}
lowids <- getPMIDsByKeyWords(journal = lows[1], dFrom = yr, dTo = yr, n = 10000, https = TRUE)
for(i in 2:length(lows)){
  lowids <- c(lowids,getPMIDsByKeyWords(journal = lows[i], dFrom = yr, dTo = yr, n = 10000, https = TRUE))
  Sys.sleep(0.5)
}

highabstracts <- getAbstracts(highids[1])
for (j in 2:length(highids)){
  Sys.sleep(0.25)
  beep_on_error(tryCatch({
    highabstracts <- c(highabstracts,getAbstracts(highids[j]))
  }, error = function(e) {
    Sys.sleep(0.75)
    highabstracts <- c(highabstracts,getAbstracts(highids[j]))
  }), sound = 1)
  #highabstracts <- c(highabstracts,getAbstracts(highids[j]))
  #Sys.sleep(0.2)
  print(j)
  print(Sys.time())
}
beep("complete")

highabstracts <- addpotents(highabstracts)
beep("complete")

lowabstracts <- getAbstracts(lowids[1])
for (j in 2:length(lowids)){
  Sys.sleep(0.25)
  beep_on_error(tryCatch({
    lowabstracts <- c(lowabstracts,getAbstracts(lowids[j]))
  }, error = function(e) {
    Sys.sleep(0.75)
    lowabstracts <- c(lowabstracts,getAbstracts(lowids[j]))
  }), sound = 1)
  #lowabstracts <- c(lowabstracts,getAbstracts(lowids[j]))
  #Sys.sleep(0.2)
  print(j)
  print(Sys.time())
}
beep("fanfare")

abcorp <- Corpus(VectorSource(highabstracts))

abmap <- tm_map(abcorp, PlainTextDocument)
abmap <- tm_map(abmap, removePunctuation)
abmap <- tm_map(abmap, removeWords, c(stopwords('english'),scienceywords))
#abmap <- tm_map(abmap, stemDocument)
# I personally don't like to stem the words

termy <- clean_corpus(abmap)
termy <- breakwords(termy)
term_dtm <- DocumentTermMatrix(termy)
# According to https://www.tidytextmining.com/dtm.html 'Converting to and from...' this is the step
# where we need to convert from corpus into text data using the 'tidy' function

high_words <- tidy(term_dtm)
high_words$tier <- 'High'
high_words$total <- sum(high_words$count)

abcorp <- Corpus(VectorSource(lowabstracts))

abmap <- tm_map(abcorp, PlainTextDocument)
abmap <- tm_map(abmap, removePunctuation)
abmap <- tm_map(abmap, removeWords, stopwords('english'))

termy <- clean_corpus(abmap)
termy <- breakwords(termy)
term_dtm <- DocumentTermMatrix(termy)

low_words <- tidy(term_dtm)
low_words$tier <- 'Low'
low_words$total <- sum(low_words$count)

abstracts <- bind_rows(low_words, high_words)

frequency <- abstracts %>% 
  group_by(tier) %>% 
  count(term, sort = TRUE) %>% 
  left_join(abstracts %>% 
              group_by(tier) %>% 
              summarise(total = n())) %>%
  mutate(freq = n/total)

frequency <- frequency %>% 
  select(tier, term, freq) %>% 
  spread(tier, freq) %>%
  arrange(Low, High)

word_ratios <- abstracts %>%
  count(term, tier) %>%
  group_by(term) %>%
  filter(sum(n) >= 10) %>%
  ungroup() %>%
  spread(tier, n, fill = 0) %>%
  mutate_if(is.numeric, funs((. + 1) / (sum(.) + 1))) %>%
  mutate(logratio = log(High / Low)) %>%
  arrange(desc(logratio))

nam <- paste(as.character(yr),'Contrasts.png')
png(nam, width = 900, height = 450)

word_ratios %>%
  group_by(logratio < 0) %>%
  top_n(15, abs(logratio)) %>%
  ungroup() %>%
  mutate(term = reorder(term, logratio)) %>%
  ggplot(aes(term, logratio, fill = logratio < 0)) +
  geom_col(show.legend = FALSE) +
  coord_flip() +
  ylab("log odds ratio (High/Med)") +
  scale_fill_discrete(name = "", labels = c("High", "Med")) +
  theme_gray(base_size=12*(80/60))

dev.off()

# to view and weed out nonsense words, check dump then add them to sciencywords
dump <- word_ratios %>% group_by(logratio < 0) %>% top_n(50, abs(logratio))

# to look up the odds ratio of a specific term
# word_ratios[match('optogenetics',word_ratios$term),]

nam <- paste(as.character(yr),'Data.RData')
save.image(file = nam)
```
