## Supplementary material for "Corpus Colossal: A Bibliometric Analysis of Neuroscience Abstracts and Impact Factor": Analysis Code: Species and Neurotransmitter Contrasts.R

```
# library(beepr)
# library(tm)
# library(PubMedWordcloud)
# library(RColorBrewer)
# library(SnowballC)
# library(tidytext)
# library(dplyr)
# library(tidyr)
# library(ggplot2)
# library(stringr)
# 
# load('Master Data.RData')

word_ratios <- abstracts %>%
  count(term, tier) %>%
  group_by(term) %>%
  filter(sum(n) >= 50) %>%
  filter(!str_detect(term, "copyright")) %>%
  ungroup() %>%
  spread(tier, n, fill = 0) %>%
  mutate(instance = as.character(High + Low)) %>%
  mutate_if(is.numeric, funs((. + 1) / (sum(.) + 1))) %>%
  mutate(logratio = log(High / Low)) %>%
  arrange(desc(logratio))
word_ratios$instance <- as.numeric(word_ratios$instance)

# let's do a species comparison
species <- c("drosophila" , "elegans" , "rats" , "mice" , "men" , "women" , "hamsters" , "vole" , "primate" , "lizard" , "zebrafish")
tframe <-data.frame(species)
for (i in 1:length(species)){
  tframe$ratio[i]<- word_ratios[match(species[i],word_ratios$term),]$logratio
  tframe$instance[i] <- word_ratios[match(species[i],word_ratios$term),]$instance
}
tframe <- tframe[complete.cases(tframe), ]

tframe <- transform(tframe, species=reorder(species, ratio) ) 

e <- ggplot(tframe, aes(x = species, y = ratio)) + ylab("Log odds ratio (Med/High)") + xlab("Study Organism") + theme_classic()
e <- e + geom_segment(aes(x = species, y = -1.7, xend = species, yend = ratio), linetype = "dashed", colour = "gray")
e <- e + geom_point(aes(fill=ratio, size=instance), colour="black",shape=21,show.legend=FALSE, position=position_dodge(.75)) + scale_size(range = c(2, 10))
e <- e + theme(text = element_text(size=14, face="bold"), axis.line = element_line(color="black", size=1, linetype="solid"))

print(e)


# now let's do a neurotransmitter comparison
NTs <- c("dopamine" , "serotonin" , "gaba" , "glutamate" , "acetylcholine" ,"epinephrine", "norepinephrine",  "endorphins" , "oxytocin" , "vasopressin" , "testosterone" , "estrogen", "estradiol","progesterone")
tframe <-data.frame(NTs)
for (i in 1:length(NTs)){
  tframe$ratio[i]<- word_ratios[match(NTs[i],word_ratios$term),]$logratio
  tframe$instance[i] <- word_ratios[match(NTs[i],word_ratios$term),]$instance
}
tframe <- tframe[complete.cases(tframe), ]

tframe <- transform(tframe, NTs=reorder(NTs, ratio) ) 

e <- ggplot(tframe, aes(x = NTs, y = ratio)) + ylab("Log odds ratio (Med/High)") + xlab("Neurotransmitter") + theme_classic()
e <- e + geom_segment(aes(x = NTs, y = -1.5, xend = NTs, yend = ratio), linetype = "dashed", colour = "gray")
e <- e + geom_point(aes(fill=ratio, size=instance), colour="black",shape=21,show.legend=FALSE, position=position_dodge(.75)) + scale_size(range = c(3, 12))
e <- e + theme(text = element_text(size=14, face="bold"), axis.line = element_line(color="black", size=1, linetype="solid"),axis.text.x = element_text(angle = 30, hjust = 1))

print(e)


# now let's do a methods comparison
Methods <- c("fmri" , "dti" , "pet" , "microdialysis" , "optogenetic", "rtpcr", "proteomic")
Methods <- c(Methods, "behavior", "singlecell","calcium","meg","eeg")
Methods <- c(Methods, "restingstate", "knockout", "transcranial","gwas","transcriptome", "spiking")
Methods <- c(Methods, "agonist", "antagonist", "vivo", "vitro")

Names <- c("fMRI", "DTI", "PET","microdialysis" , "optogenetic", "rt-PCR", "proteomic")
Names <- c(Names, "behavior", "single-cell","calcium","MEG","EEG")
Names <- c(Names, "resting-state", "knockout", "transcranial","GWAS","transcriptome", "spiking")
Names <- c(Names, "agonist", "antagonist", "vivo", "vitro")

Tried <- c("bold","chemogenetics", "chemogenetically","dreadd", "dreadds","proteomics","optogenetically")
Tried <- c(Tried,"transcriptomic","knockdown","voltametry","rnaseq")

tframe <-cbind(Methods,Names)
tframe <-data.frame(tframe)
for (i in 1:length(Methods)){
  tframe$ratio[i]<- word_ratios[match(Methods[i],word_ratios$term),]$logratio
  tframe$instance[i] <- word_ratios[match(Methods[i],word_ratios$term),]$instance
}
tframe <- tframe[complete.cases(tframe), ]

tframe <- tframe[order(tframe$ratio),] 

tframe <- transform(tframe, Methods=reorder(Methods, ratio) )

e <- ggplot(tframe, aes(x =reorder(Methods, ratio), y = ratio)) + ylab("Log odds ratio (Med/High)") + xlab("Approach") + theme_classic()
e <- e + scale_y_continuous(limits = c(-3.1, 2.2)) + geom_segment(aes(x = Methods, y = -3.1, xend = Methods, yend = ratio), linetype = "dashed", color = "gray")
e <- e + geom_point(aes(fill=ratio*-1, size=instance),shape=21,show.legend=FALSE, position=position_dodge(.75)) + scale_fill_gradientn(colours = terrain.colors(2)) + scale_size(range = c(3, 12))
e <- e + theme(text = element_text(size=14, face="bold"), axis.line = element_line(color="black", size=1, linetype="solid"),axis.text.x = element_text(angle = 30, hjust = 1))
e <- e + scale_x_discrete(labels= tframe$Names)

print(e)


# now let's do brain regions

regions <- c("cerebellum", "cortex", "hippocampus", "amygdala", "hypothalamus", "thalamus", "accumbens", "striatum", "tegmental")
regions <- c(regions, "hindbrain", "pons", "medulla", "prefrontal", "orbitofrontal", "raphe", "insula", "spinal", "paraventricular", "dentate")
regions <- c(regions, "septum", "terminalis", "periaqueductal", "preoptic")

Names <- c("Cerebellum", "Cortex", "Hippocampus", "Amygdala", "Hypothalamus", "Thalamus", "Nuc. Accumbens", "Striatum", "Tegmental")
Names <- c(Names, "Hindbrain", "Pons", "Medulla", "Prefrontal", "Orbitofrontal", "Raphe", "Insula", "Spinal", "Paraventricular", "Dentate")
Names <- c(Names, "Septum", "Terminalis", "Periaqueductal", "Preoptic")

tframe <-cbind(regions,Names)
tframe <-data.frame(tframe)
for (i in 1:length(regions)){
  tframe$ratio[i]<- word_ratios[match(regions[i],word_ratios$term),]$logratio
  tframe$instance[i] <- word_ratios[match(regions[i],word_ratios$term),]$instance
}
tframe <- tframe[complete.cases(tframe), ]

tframe <- tframe[order(tframe$ratio),] 

tframe <- transform(tframe, regions=reorder(regions, ratio) )

e <- ggplot(tframe, aes(x =reorder(regions, ratio), y = ratio)) + ylab("Log odds ratio (Med/High)") + xlab("Brain Region") + theme_classic()
e <- e + scale_y_continuous(limits = c(-0.8, 0.5)) + geom_segment(aes(x = regions, y = -0.8, xend = regions, yend = ratio), color = "gray")
e <- e + geom_point(aes(fill=ratio*-1, size=instance),shape=21,show.legend=FALSE, position=position_dodge(.75)) + scale_fill_gradient(low="mediumpurple4",high="lightpink") + scale_size(range = c(3, 12))
e <- e + theme(text = element_text(size=14, face="bold"), axis.line = element_line(color="black", size=1, linetype="solid"),axis.text.x = element_text(angle = 30, hjust = 1))
e <- e + scale_x_discrete(labels= tframe$Names)

print(e)
```
