## Supplementary material for "Corpus Colossal: A Bibliometric Analysis of Neuroscience Abstracts and Impact Factor": Analysis Code: Theme Contrasts.R

```
# library(beepr)
# library(tm)
# library(PubMedWordcloud)
# library(RColorBrewer)
# library(SnowballC)
# library(tidytext)
# library(dplyr)
# library(tidyr)
# library(ggplot2)
# library(stringr)
# 
# load('Master Data.RData')

word_ratios <- abstracts %>%
  count(term, tier) %>%
  group_by(term) %>%
  filter(sum(n) >= 50) %>%
  filter(!str_detect(term, "copyright")) %>%
  ungroup() %>%
  spread(tier, n, fill = 0) %>%
  mutate(instance = as.character(High + Low)) %>%
  mutate_if(is.numeric, funs((. + 1) / (sum(.) + 1))) %>%
  mutate(logratio = log(High / Low)) %>%
  arrange(desc(logratio))
word_ratios$instance <- as.numeric(word_ratios$instance)

# let's make theme clusters and have them compete!

topics <- c("neuroimmunology", "inflammation", "cytokine", "cytokines", "interleukin", "proinflammatory", "neuroinflammation")
tframe <-data.frame(topics)
tframe$theme <- 'neuro- \n immunology'
for (i in 1:length(topics)){
  tframe$ratio[i]<- word_ratios[match(topics[i],word_ratios$term),]$logratio
}
tframe$rank <- median(tframe$ratio, na.rm=TRUE)
clusters <- tframe

topics <- c("neurogenesis", "stem", "progenitor", "progenitors")
tframe <-data.frame(topics)
tframe$theme <- 'neurogenesis'
for (i in 1:length(topics)){
  tframe$ratio[i]<- word_ratios[match(topics[i],word_ratios$term),]$logratio
}
tframe$rank <- median(tframe$ratio, na.rm=TRUE)
clusters <- rbind(tframe,clusters)

topics <- c("comparative", "ethological", "neuroethology", "species", "evolutionary","ethology")
tframe <-data.frame(topics)
tframe$theme <- 'comparative'
for (i in 1:length(topics)){
  tframe$ratio[i]<- word_ratios[match(topics[i],word_ratios$term),]$logratio
}
#clusters <- rbind(tframe,clusters)
# not included due to half NAs

topics <- c("epigenetics", "epigenetic", "transgenerational", "methylation", "histone", "cpg")
tframe <-data.frame(topics)
tframe$theme <- 'epigenetics'
for (i in 1:length(topics)){
  tframe$ratio[i]<- word_ratios[match(topics[i],word_ratios$term),]$logratio
}
tframe$rank <- median(tframe$ratio, na.rm=TRUE)
clusters <- rbind(tframe,clusters)

topics <- c("gut", "microbiome", "microbiota", "germfree", "probiotic","intestine", "intestinal", "gutbrainaxis", "microbial", "microbe")
tframe <-data.frame(topics)
tframe$theme <- 'microbiome'
for (i in 1:length(topics)){
  tframe$ratio[i]<- word_ratios[match(topics[i],word_ratios$term),]$logratio
}
tframe$rank <- median(tframe$ratio, na.rm=TRUE)
clusters <- rbind(tframe,clusters)

topics <- c("stress", "corticosteriod", "cortisol", "corticosterone", "glucocorticoid", "hpa", "hpaaxis", "adrenal")
tframe <-data.frame(topics)
tframe$theme <- 'stress'
for (i in 1:length(topics)){
  tframe$ratio[i]<- word_ratios[match(topics[i],word_ratios$term),]$logratio
}
tframe$rank <- median(tframe$ratio, na.rm=TRUE)
clusters <- rbind(tframe,clusters)

topics <- c("social", "maternal", "empathy", "prosocial", "affiliative")
tframe <-data.frame(topics)
tframe$theme <- 'social'
for (i in 1:length(topics)){
  tframe$ratio[i]<- word_ratios[match(topics[i],word_ratios$term),]$logratio
}
tframe$rank <- median(tframe$ratio, na.rm=TRUE)
clusters <- rbind(tframe,clusters)

topics <- c("alzheimers", "tau", "amyloid", "plaque", "apoe")
tframe <-data.frame(topics)
tframe$theme <- 'alzheimers'
for (i in 1:length(topics)){
  tframe$ratio[i]<- word_ratios[match(topics[i],word_ratios$term),]$logratio
}
tframe$rank <- median(tframe$ratio, na.rm=TRUE)
clusters <- rbind(tframe,clusters)

topics <- c("drug", "drugs", "abuse", "reward", "addiction")
tframe <-data.frame(topics)
tframe$theme <- 'drug \n abuse'
for (i in 1:length(topics)){
  tframe$ratio[i]<- word_ratios[match(topics[i],word_ratios$term),]$logratio
}
tframe$rank <- median(tframe$ratio, na.rm=TRUE)
clusters <- rbind(tframe,clusters)

topics <- c("depression", "depressive", "anxiety", "mdd")
tframe <-data.frame(topics)
tframe$theme <- 'depression'
for (i in 1:length(topics)){
  tframe$ratio[i]<- word_ratios[match(topics[i],word_ratios$term),]$logratio
}
tframe$rank <- median(tframe$ratio, na.rm=TRUE)
clusters <- rbind(tframe,clusters)

topics <- c("learning","memory","retrieval","retention","habituation","conditioned","extinction","retrieval")
tframe <-data.frame(topics)
tframe$theme <- 'learning \n memory'
for (i in 1:length(topics)){
  tframe$ratio[i]<- word_ratios[match(topics[i],word_ratios$term),]$logratio
}
tframe$rank <- median(tframe$ratio, na.rm=TRUE)
clusters <- rbind(tframe,clusters)

clusters <- transform(clusters, theme=reorder(theme, rank) ) 

e <- ggplot(clusters, aes(x = theme, y = ratio)) + ylab("Log odds ratio (Med/High)") + xlab("Theme") + theme_classic()
e <- e + geom_boxplot(aes(fill = theme), size = 1)  
#e <- e + scale_fill_manual(values = c("#BFBFBF", "#9C71AB"))
# and now the dots
e <- e + geom_point(aes(fill=theme), colour="black",shape=21,size=4,show.legend=FALSE, position=position_dodge(.75))
#e <- e + scale_color_manual(values = c("#BFBFBF", "#9C71AB"))
# set up the text:
e <- e + theme(text = element_text(size=14, face="bold"), axis.line = element_line(color="black", size=1, linetype="solid"),axis.text.x = element_text(angle = 30, hjust = 1))
print(e)
```
