## Supplementary material for "Corpus Colossal: A Bibliometric Analysis of Neuroscience Abstracts and Impact Factor": Analysis Code: Trends by Year.R

```
rm(list = ls())
setwd("D:/Box Sync/Research/Meta Review")

#the original formulation left out instance, so make sure that gets included
includeinst <- function(highabstracts)
{
  word_ratios <- abstracts %>%
    count(term, tier) %>%
    group_by(term) %>%
    filter(sum(n) >= 50) %>%
    filter(!str_detect(term, "copyright")) %>%
    ungroup() %>%
    spread(tier, n, fill = 0) %>%
    mutate(instance = as.character(High + Low)) %>%
    mutate(numhigh = as.character(High)) %>%
    mutate(numlow = as.character(Low)) %>%
    mutate_if(is.numeric, funs((. + 1) / (sum(.) + 1))) %>%
    mutate(logratio = log(High / Low)) %>%
    arrange(desc(logratio))
  word_ratios$instance <- as.numeric(word_ratios$instance)
  return(word_ratios)
}


load("D:/Box Sync/Research/Meta Review/2014 Data.RData")
word_ratios <- includeinst(word_ratios)
names(word_ratios) <- c("term", paste("High",yr,sep=""), paste("Low",yr,sep=""), paste("instance",yr,sep=""), paste("numhigh",yr,sep=""), paste("numlow",yr,sep=""), paste("logratio",yr,sep=""))
trends <- word_ratios

load("D:/Box Sync/Research/Meta Review/2015 Data.RData")
word_ratios <- includeinst(word_ratios)
names(word_ratios) <- c("term", paste("High",yr,sep=""), paste("Low",yr,sep=""), paste("instance",yr,sep=""), paste("numhigh",yr,sep=""), paste("numlow",yr,sep=""), paste("logratio",yr,sep=""))
trends <- merge(trends,word_ratios,all = TRUE)

load("D:/Box Sync/Research/Meta Review/2016 Data.RData")
word_ratios <- includeinst(word_ratios)
names(word_ratios) <- c("term", paste("High",yr,sep=""), paste("Low",yr,sep=""), paste("instance",yr,sep=""), paste("numhigh",yr,sep=""), paste("numlow",yr,sep=""), paste("logratio",yr,sep=""))
trends <- merge(trends,word_ratios,all = TRUE)

load("D:/Box Sync/Research/Meta Review/2017 Data.RData")
word_ratios <- includeinst(word_ratios)
names(word_ratios) <- c("term", paste("High",yr,sep=""), paste("Low",yr,sep=""), paste("instance",yr,sep=""), paste("numhigh",yr,sep=""), paste("numlow",yr,sep=""), paste("logratio",yr,sep=""))
trends <- merge(trends,word_ratios,all = TRUE)

load("D:/Box Sync/Research/Meta Review/2018 Data.RData")
word_ratios <- includeinst(word_ratios)
names(word_ratios) <- c("term", paste("High",yr,sep=""), paste("Low",yr,sep=""), paste("instance",yr,sep=""), paste("numhigh",yr,sep=""), paste("numlow",yr,sep=""), paste("logratio",yr,sep=""))
trends <- merge(trends,word_ratios,all = TRUE)

sortee <- order(names(trends))
sortee <- c(1,sortee[1:length(sortee)-1])
trends <- trends[, sortee]

write.csv(trends,"trends.csv",na="")
# now calcutlate slope of change between years in excel like the caveman you are and re-load
trends <- read.csv("trends.csv")

for (i in 1:length(scienceywords)){
  trends<- filter(trends, !grepl(scienceywords[i], term))
}


trends %>%
  group_by(ratioslope < 0) %>%
  top_n(25, abs(ratioslope)) %>%
  ungroup() %>%
  mutate(term = reorder(term, ratioslope)) %>%
  ggplot(aes(term, ratioslope, fill = ratioslope < 0)) +
  geom_col(show.legend = FALSE) +
  coord_flip() +
  ylab("Slope of change in log odds ratio from 2014-2018") +
  scale_fill_discrete(name = "", labels = c("High", "Med"))

trends %>%
  group_by(ratioslope < 0) %>%
  top_n(25, abs(ratioslope)) %>%
  ungroup() %>%
  mutate(term = reorder(term, ratioslope)) %>%
  ggplot(aes(term, ratioslope, fill = ratioslope < 0)) +
  geom_col(show.legend = FALSE) +
  coord_flip() +
  ylab("Slope of change in log odds ratio from 2014-2018") +
  scale_fill_discrete(name = "", labels = c("High", "Med"))

steadies <- trends
steadies %>%
  group_by(logratio2014 < 0) %>%
  top_n(25, abs(logratio2014)) %>%
  ungroup() %>%
  mutate(term = reorder(term, logratio2014)) %>%
  ggplot(aes(term, logratio2014-logratio2018, fill = logratio2018 < 0)) +
  geom_col(show.legend = FALSE) +
  coord_flip() +
  ylab("log odds ratio (High/Med)") +
  scale_fill_discrete(name = "", labels = c("High", "Med")) +
  theme_gray(base_size=12*(80/60))


# this results in a bunch of boring 'brain', 'neuron', 'cell'...
trends %>%
  group_by(highslope < 0) %>%
  top_n(25, abs(highslope)) %>%
  ungroup() %>%
  mutate(term = reorder(term, highslope)) %>%
  ggplot(aes(term, highslope, fill = highslope < 0)) +
  geom_col(show.legend = FALSE) +
  coord_flip() +
  ylab("Slope of Increase in # of high-impact pubs") +
  scale_fill_discrete(name = "", labels = c("High", "Med"))
```
